# Collective invasion of the basement membrane in breast cancer driven by forces from cell volume expansion and local contractility

**DOI:** 10.1101/2022.07.28.501930

**Authors:** Julie Chang, Aashrith Saraswathibhatla, Zhaoqiang Song, Sushama Varma, Colline Sanchez, Sucheta Srivastava, Katherine Liu, Michael C. Bassik, M. Peter Marinkovich, Louis Hodgson, Vivek Shenoy, Robert B. West, Ovijit Chaudhuri

## Abstract

Breast cancer becomes invasive when carcinoma cells collectively invade through the basement membrane (BM), a nanoporous layer of matrix that physically separates the primary tumor from the stroma, in a first step towards metastasis. Single cells can invade through nanoporous three-dimensional (3D) matrices via protease-mediated degradation or, when the matrix exhibits sufficient mechanical plasticity, force-mediated widening of pores. However, how cells invade collectively through physiological BM layers in cancer remains unclear. Here, we developed a 3D *in vitro* model of collective invasion of the BM during breast cancer. We show that cells utilize both proteases and forces to breach the BM. Forces are generated from a combination of global cell volume expansion that stretch the BM with local contractile forces that act in the plane of the BM to breach it, allowing invasion. These results uncover a mechanism by which cells collectively interact to overcome a critical barrier to metastasis.

## Main

The initial invasion of carcinoma cells through the basement membrane (BM) and into the stromal matrix is a necessary first step to metastasis.^1^ The BM is a dense, nanoporous meshwork of extracellular matrix (ECM) proteins composed of primarily laminin and collagen-IV that separates epithelial cells from the surrounding stromal matrix and acts as a physical barrier to cell invasion.^2, 3^ In pre-invasive ductal carcinoma in situ (DCIS), mammary epithelial cells undergo aberrant proliferation, but remain confined within the BM, and this pre-invasive condition is considered largely treatable with low mortality.^4, 5^ However, once the carcinoma cells invade through the BM and into the stroma, the tumor becomes invasive breast carcinoma (IBC) and can then undergo metastasis, which results in increased mortality.^6^ Initial invasion of the BM is primarily collective in nature, driven by groups of cells as opposed to individual cells.^7^ Enhanced mammographic density is one of the strongest and most consistent risk factors for breast cancer progression, and, correspondingly, increased ECM stiffness, associated with enhanced mammographic density, is thought to play a key role in promoting invasive breast cancer progression (Fig. 1a).^8^ Therefore, the BM and the stiffness of the surrounding ECM are important gatekeepers for tumor progression from a benign into an invasive and malignant phenotype.

**Figure 1.**
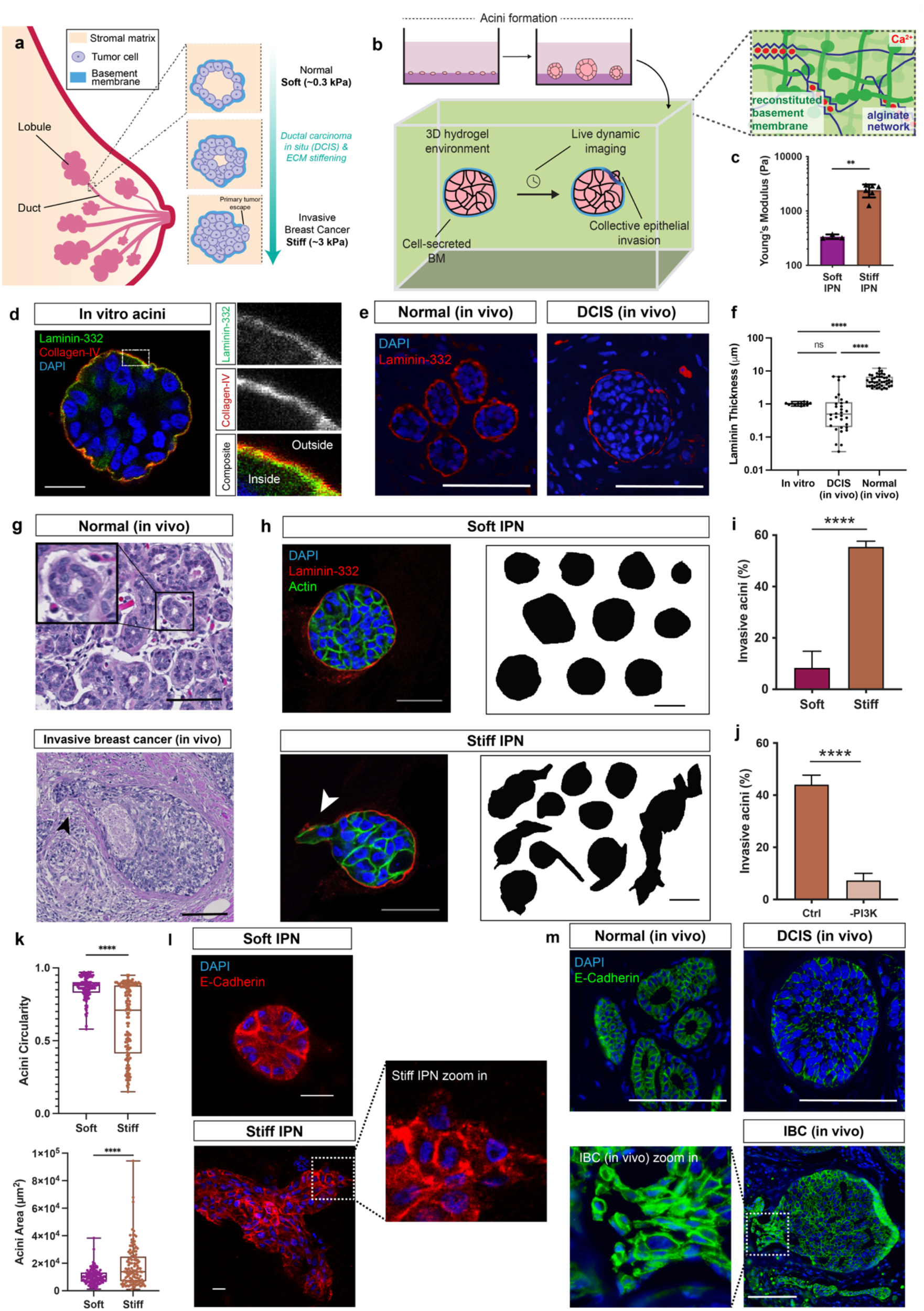
Increased stiffness induces collective invasion through endogenous basement membrane in 3D culture model of invasive breast carcinoma. **a**, Schematic of breast carcinoma progression. **b**, Schematic of 3D *in vitro* model of breast carcinoma progression, in which pre-formed organotypic MCF10A acini are encapsulated into soft or stiff alginate-rBM IPNs. **c**, Young’s modulus of soft and stiff alginate-rBM IPNs. \*\**p* < 0.01, unpaired t-test, *n =* 3 – 6 hydrogels. **d**, Collagen-IV (red), laminin-332 (green), and 4,6-diamidino-2-phenylindole (DAPI) (blue) staining of endogenous basement membrane structure in MCF10A acini. Scale bar, 20 μm. **e**, Laminin-332 (red) and DAPI (blue) in normal and DCIS *in vivo* breast tissue samples. Scale bar, 100 μm. **f**, Laminin thickness of *in vitro* acini and *in vivo* DCIS and normal samples. n = 12 *in vitro* samples, n = 32 *in vivo* DCIS samples, n = 45 *in vivo* normal samples. ns = not statistically significant. ****p < 0.0001, ANOVA. **g**, H&E images of normal (top) and invasive breast cancer (bottom) *in vivo* breast tissue samples. Black arrow points to collectively invading cells. Scale bar, 100 μm. **h**, Laminin-332 (red), actin (green), and DAPI (blue) of MCF10A acini in soft and stiff IPNs. Traces of acini in soft and stiff IPNs. Scale bar, 50 μm. **i**, Percentage of invasive acini in soft and stiff IPNs. \*\*\*\**p* < 0.0001, Welch’s t-test. **j**, Percentage of invasive acini in stiff IPNs with control or PI3K inhibitor. \*\*\*\**p* < 0.0001, unpaired t-test, *n =* >30 acini in 3 – 6 hydrogels. **k**, Quantification of acini circularity and acini area in soft and stiff IPNs. \*\*\*\**p* < 0.0001, Welch’s t-test. **l**, Actin (green) and E-cadherin (red) staining in soft and stiff IPNs. Scale bars, 20 μm. **m**, E-cadherin (green) and DAPI (blue) in normal, DCIS, and invasive breast cancer *in vivo* breast tissue samples. Scale bar, 100um.

In principle, the BM can be breached by carcinoma cells acting through either protease-dependent, chemical modes, or force driven, protease-independent modes. It has been long established that cells can secrete matrix metalloproteinases (MMPs) to chemically degrade the BM and create holes that enable cancer cell invasion into the surrounding stromal matrix and eventually, metastasize.^9, 10^ Specifically, individual cells can extend actin-rich, invasive protrusions known as invadopodia, which secrete MMPs and also express membrane anchored MMPs.^11^ Complementarily, recent studies reveal that invadopodia and cells as a whole can exert forces, without the aid of proteolytic degradation, to physically deform, remodel, and move through even nanoporous ECM if the ECM exhibits sufficient mechanical plasticity (i.e. malleability).^12–14^ Plasticity is defined as the irreversible deformation that occurs after a mechanical load is applied.^15^ Increased stiffness of the BM itself and the stromal matrix has been shown to promote these invasive behaviors and subsequently, metastasis.^16, 17^ However, most studies of BM invasion primarily focus on non-cancer cells or models using exogenous BM or reconstituted BM matrix. Thus, how cancer cells use proteases and/or forces to breach endogenous BM are undescribed.

An important feature of initial invasion through the BM is that cells migrate and invade collectively, rather than individually. In collective groups, cells attach to each other via cell-cell junctions, such as adherens junctions, which coordinate collective movements.^18^ While the loss of E-cadherin, a widely studied adherens junctions component, has been shown to promote cell invasion, this loss is likely to be more relevant to single cell dissemination.^19^ E-cadherin is expressed in many invasive forms of breast carcinoma, suggesting that collective, rather than single, cell migration is the dominant mode of migration.^20^ Collective invasion has been studied extensively via hydrogels composed of type 1 collagen, the primary component of the stromal matrix that cells migrate through after invading through the BM. Key findings from these models demonstrate that confinement promotes collective invasion and leader cells expressing basal epithelial genes drive invasion.^21, 22^ Studies of collective invasion in a BM rich matrix model using increased stiffness to induce invasion by normal mammary epithelium showed that epithelial cells undergo softening and swelling during invasion.^23^ However, these studies do not focus on the initial invasion through the BM, in which cells directly interact with and invade through the BM as a thin barrier before migrating into the stromal matrix. What, if any, is the functional significance of collective interactions for BM invasion remains unclear.

Here we studied the dynamics of collective invasion through endogenous BM in a 3D *in vitro* model of breast cancer. Using an *in vitro* model of stiffness induced invasion of mammary epithelial cells through endogenous BM, we studied the mechanisms by which cells collectively breach through the BM (Fig. 1a). These data reveal that cells use both protease and force-driven modes of invasion through the BM. Mechanistically, we find that cells synergistically use both global cell volume expansion and localized, actomyosin contractility to exert tangential forces on the BM and create openings for cells to migrate through during invasion.

## Results

### 3D in vitro model of breast carcinoma invasion

We designed a three-dimensional (3D) culture model using hydrogels to study collective mammary epithelial cell invasion through endogenous BM (Fig 1a, b). Single MCF10A mammary epithelial cells were cultured on a layer of rBM matrix, during which they proliferated and self-assembled into 3D organotypic acini, in which the cells exhibit key features observed in normal mammary epithelial cells in ducts.^24^ Specifically, these acini exhibit apicobasal polarity, lumen formation, and secrete a layer of endogenous BM on the periphery. The acini were harvested and encapsulated into hydrogels comprised of interpenetrating networks (IPNs) of rBM and alginate. Through the addition of calcium ions, IPN stiffness can be modulated, without changing protein concentration, matrix architecture, or pore size.^25^ Note that calcium used for crosslinking itself does not impact the phenotype of these cells.^25^ IPNs of two stiffnesses were made: “soft” IPNs that mimicked the stiffness of healthy tissue had an elastic modulus 0.3 kPa and “stiff” IPNs that mimicked the increased stiffness of tumor tissue had an elastic modulus of 2.5 kPa (Fig. 1c).^26, 27^ These IPNs exhibited similar degrees of mechanical plasticity in creep-recovery tests and viscoelasticity, as indicated by the loss tangents, in rheological analysis (Fig. S1). Overall, we developed IPNs of different stiffness but maintaining similar mechanical plasticity and viscoelastic behaviors.

Next, we evaluated whether the endogenous BM secreted by MCF10A cells resembled *in vivo* BM by analyzing BM composition and thickness. The endogenous BM secreted by MCF10A cells contained laminin-332 and collagen-IV, the primary components of the BM (Fig. 1d). Laminin-332 interfaced with cells on the inner layer of the BM, while collagen-IV interfaced with the IPN on the outer layer of the BM, mimicking the *in vivo* localization of laminin and collagen-IV in the BM (Fig. 1d).^3^ Moreover, the thickness of the laminin layer in *in vitro* acini at day 7 matched the thickness of the laminin layer in *in vivo* DCIS breast tissue samples; the thickness of laminin in normal breast tissue samples were significantly thicker than laminin in DCIS samples (Fig. 1e, f). While the BM separates the acini from the IPN, previous work indicates that cells can sense IPN stiffness through endogenous BM,^28^ likely due to the mechanical tethering between the collagen-IV and laminin in both the endogenous BM and the rBM in the IPN. Therefore, the BM formed by the acini serves as a model of BM in pre-invasive breast cancer (DCIS).

We then evaluated the impact of increased IPN stiffness on cell invasion and the ability of this *in vitro* model to recapitulate key features of human breast carcinoma progression. In human breast tissue samples, normal ducts appeared round, while ducts in invasive breast carcinoma exhibited a jagged and irregular morphology (Fig. 1g). These two states were captured by the stable round acini in soft IPNs and the invasive acini observed in the stiff IPNs, respectively (Fig. 1h). Moreover, BM around the soft IPNs remained intact, while cells breached the BM in the stiff IPNs. Overall, more acini became invasive in stiff IPNs compared to soft IPNs after 3 days of encapsulation in IPNs (Fig. 1i), indicating that increased ECM stiffness leads to the invasion of mammary epithelium by breaching the endogenous BM. To test if the cells in the invasive acini exhibited malignant phenotypes resembling that of invasive ductal carcinoma, we inhibited PI3K pathway which is often activated during hyperplasia, prior to breast cancer progression.^29^ Upon PI3K inhibition in acini embedded in stiff IPNs, the percentage of the invasive acini significantly decreased (Fig. 1j), suggesting that invasive phenotype observed here is induced through at least some of the same signaling pathways characteristic of the malignant phenotype of invasive ductal carcinoma. Further quantification of cell invasion metrics demonstrated lower acinar circularity and higher acinar area in stiff IPNs compared to soft IPNs, supporting the finding of increased collective invasion in stiff IPNs (Fig. 1k). Cell-cell junction protein E-cadherin was present in both non-invasive acini in soft IPNs and invasive acini in stiff IPNs, demonstrating E-cadherin positive collective invasion in this model, matching what is observed in vivo in invasive breast carcinoma (Fig. 1l, 1m). In summary, we developed a 3D *in vitro* model of stiffness induced collective invasion of mammary epithelial cells through endogenous BM that captures key features of invasive breast carcinoma.

### Protease inhibitors partially, but do not fully, inhibit cell invasion

After establishing a 3D *in vitro* model of invasive breast carcinoma, we sought to elucidate whether cells use protease-dependent and/or protease-independent mechanisms to invade through the BM. As greater occurrences of BM breaching were observed in acini encapsulated in stiff IPNs compared to soft IPNs, subsequent analyses of BM invasion were performed in acini encapsulated in stiff IPNs (Fig. S2). To ascertain the role of proteases in collective cell invasion, we added broad-spectrum matrix metalloproteinase (MMP) inhibitors, including GM6001, marimastat, and a previously used protease inhibitor cocktail (PIC), to the cell culture media in stiff IPNs.^30^ GM6001 and marimastat significantly reduced cell invasion, while PIC did not significantly reduce cell invasion (Fig. 2a, b). Still, cell invasion occurred when protease inhibitors were added, indicating that protease inhibitors partially, but do not fully, inhibit cell invasion.

**Figure 2.**
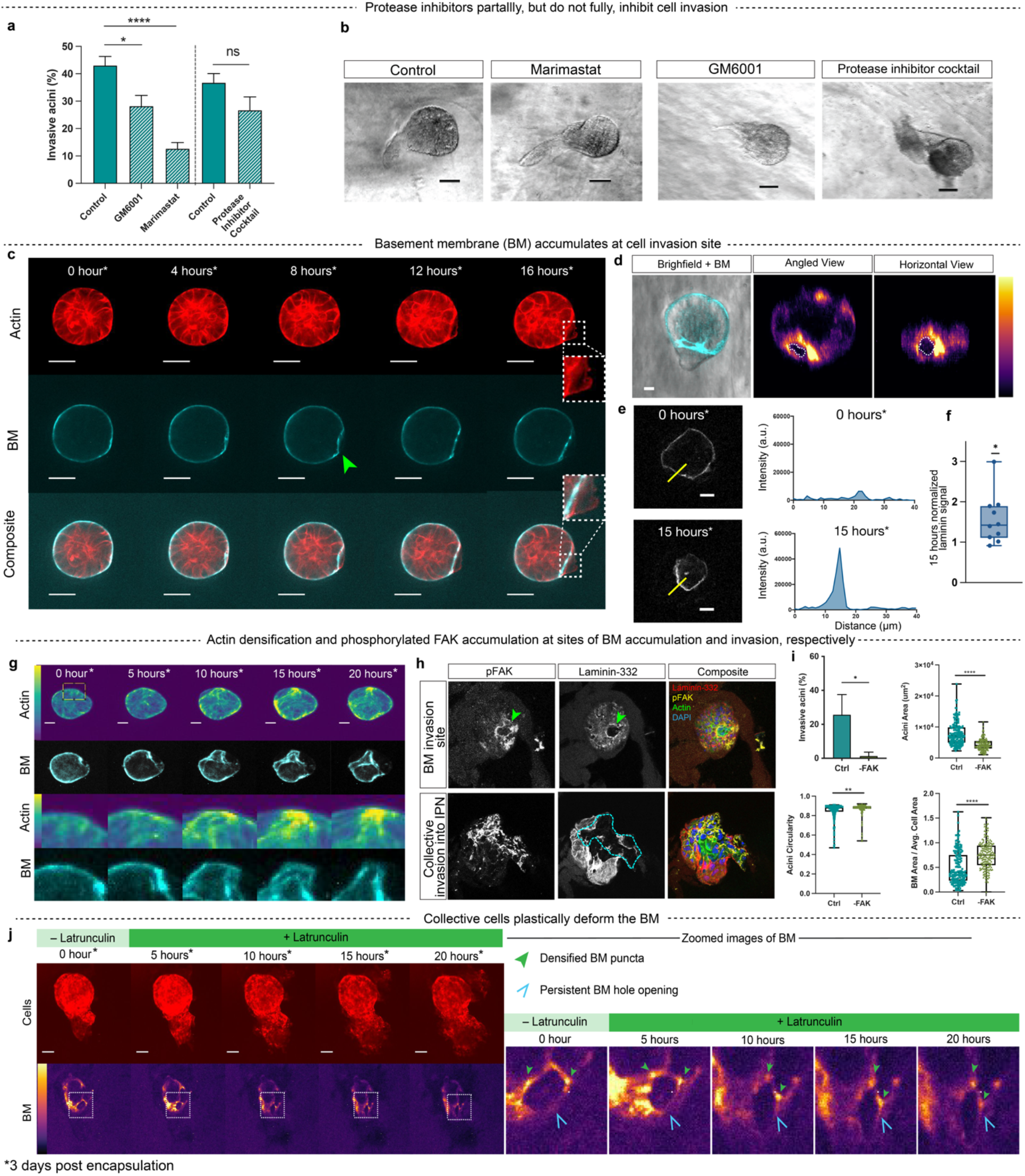
Collectively invading acini use a combination of both proteases and force to breach the basement membrane. **a,** Percentage of invasive acini with control, GM6001, marimastat, or protease cocktail inhibitor. \*\*\*\**p* < 0.0001, \**p* < 0.*05,* ns = not significant, one-way ANOVA. **b,** Representative brightfield images of invasive acini with MMP inhibitors. Scale bars, 50 μm. **c,** Time-lapse of an acini with endogenous BM exhibiting cell invasion and BM breaching. Green arrow points to BM breaching. White arrow points to cell invasion. Scale bars, 50 μm **d,** Brightfield image of invasive acini with BM in cyan. Horizontal and angled view of BM from time-lapse shown on an inferno color scale. **e,** Fluorescent images of BM in invasive acini over fifteen hours. Intensity of laminin of yellow cross-section line is plotted. Scale bar, 20 μm. **f,** Fifteen hours of normalized maximum laminin signal n = 10, * *p* < 0.05, one sample t-test. **g,** Actin signal of MCF10A-RFP LifeAct acini over twenty hours. Actin (viridis) and BM (cyan). Inset zoomed area shown in bottom row. Scale bars, 25 μm. **h,** pFAK and laminin-332 in stiff IPNs. Scale bars, 25 μm. Composite image shows pFAK (yellow), laminin-332 (red), actin (green), and nuclei (blue). Green arrows point to hole in basement membrane layer. Cyan dotted line outlines breached basement membrane. **i,** Percentage of invasive acini, cell area, cell circularity and BM area/avg. cell area in stiff IPN with control and FAK inhibitor. \*\*\*\**p* < 0.0001, \*\**p <* 0.01, Welch’s t-test. **j,** Addition of latrunculin, a potent actin depolymerizer, after cells have invaded the BM demonstrate BM plasticity. Actin (red) and BM (inferno). Scale bars, 50 μm.

### Basement membrane accumulate at cell invasion site due to cell-generated forces

Since protease inhibitors did not fully inhibit cell invasion, we explored the role of cell-exerted physical forces during BM invasion. To visualize BM movement during collective cell invasion, endogenous BM of acini was tagged with Alexafluor 488 conjugated-laminin-332 antibodies, since laminin-332 is specific to cell-secreted BM and the rBM in the IPN lacks this isoform of laminin.^31^ Within the experimental timeframe, an indiscernible amount of BM was secreted by cells after the acini are encapsulated into BM (Fig. S3). Time-lapse of BM and cell movement of acini encapsulated in stiff IPNs showed inward movement of the BM into the center of the acini while the BM was breached and cells migrated into the surrounding IPN (Fig. 2c, Video 1). Time-lapse imaging of BM and cells show an overall decrease in BM volume after cells invade through the BM and sequentially exit through the opening, thereby reducing the number of cells contained within the BM over time (Fig S4). From an angled and horizontal view of the BM, increased laminin accumulation was detected around the breaching site, showing physical densification of the BM layer around the opening where cells invaded through (Fig. 2d, e, f, Video 2). The increase of BM signal demonstrates cell-driven physical accumulation of BM, rather than a loss of BM signal that would point to protease degradation. Together, these results show that the BM invasion is associated with BM accumulation and densification, implicating cell-generated forces.

To establish whether BM densification and deformation was the result of cell-generated forces, we examined the role of actin cytoskeleton activity in BM invasion. When concurrently tracking BM movement and cell actin intensity, actin densification occurred at the site of BM densification; this accumulation of actin at the BM deformation site suggests that active forces are being generated by cells from the actin cytoskeleton around the BM (Fig. 2g, Videos 4-5). Focal adhesion kinase (FAK), a key signaling molecule that mediates actomyosin contractility through integrin-based adhesions, accumulated in its phosphorylated form at the boundaries of BM opening in invasive acini (Fig. 2h).^32^ Increased accumulation of phosphorylated FAK was also detected in invasive collective cells that have already breached the BM and invaded into the surrounding IPN (Fig. 2h). Addition of a FAK inhibitor in stiff IPNs decreased both cell invasion and BM invasion (Fig. 2i). This suggests that force transfer through integrins are associated with events of initial BM breaching, as well as post-BM breaching cell invasion. Increased BM density also suggests the possibility of mechanical remodeling of the BM due to cellular forces, mediated by plasticity of the BM. To test the role of BM mechanical plasticity in mediating BM breaching, we conducted time-lapse studies in which invading acini were exposed to latrunculin, a potent inhibitor of actin polymerization, which would be expected to release any force exerted by the cells on the BM. After addition of latrunculin, holes made in the BM remained (Fig. 2j, Videos 6-7). Moreover, high density BM puncta surrounding the hole created by collective cells also persisted after latrunculin was added, indicating plastic or irreversible deformation of the BM in response to cellular forces (Fig. 2j, green arrows). Together, our 3D *in vitro* model of invasive breast carcinoma demonstrated that collective cells use both proteases and forces to plastically deform and breach endogenous BM.

### Invadopodial protrusions do not drive BM invasion

We next sought to examine the mechanisms by which cells invade through the BM, starting with examining the role of invadopodial protrusions, which are known to drive single cell invasion.^12, 33, 34^ Invadopodial protrusions are actin rich protrusions that are hundreds of nanometers in thickness but up to tens of microns in length, display Tks5 and cortactin, are oscillatory with a lifetime on the order of hours, and deliver both proteases and forces onto the matrix to lead invasion. Comparison of collectively invading cells in MCF10A acini with invadopodia mediated invasion by single MCF10A cells and single MDA-MB-231 cells showed striking differences in the invasive behavior (Fig. 3a-c). Specifically, invadopodial protrusions were not observed in collectively invading cells in this hydrogel system; instead broader protrusions were formed, with invadopodia widths significantly narrower than the invading edge of collectively invading cells (Fig. 3d). To further test the role of invadopodia in collective invasion, Tks5 and cortactin, two key proteins involved in invadopodia protrusions, were evaluated. IHC of cortactin shows that cortactin to localize to the periphery of the invasive clusters (Fig. 3e). Next, we examined the role of Tks5, using CRISPR/Cas9 to knock out Tks5. Knockout of Tks5 in MCF10A clusters reduced invasion compared to control cells (Fig. 3f-h). These results suggest that while individual invadopodia protrusions are not observed in collective cell invasion, Tks5 and cortactin may still play a role in a different type of invasive structure.

**Figure 3.**
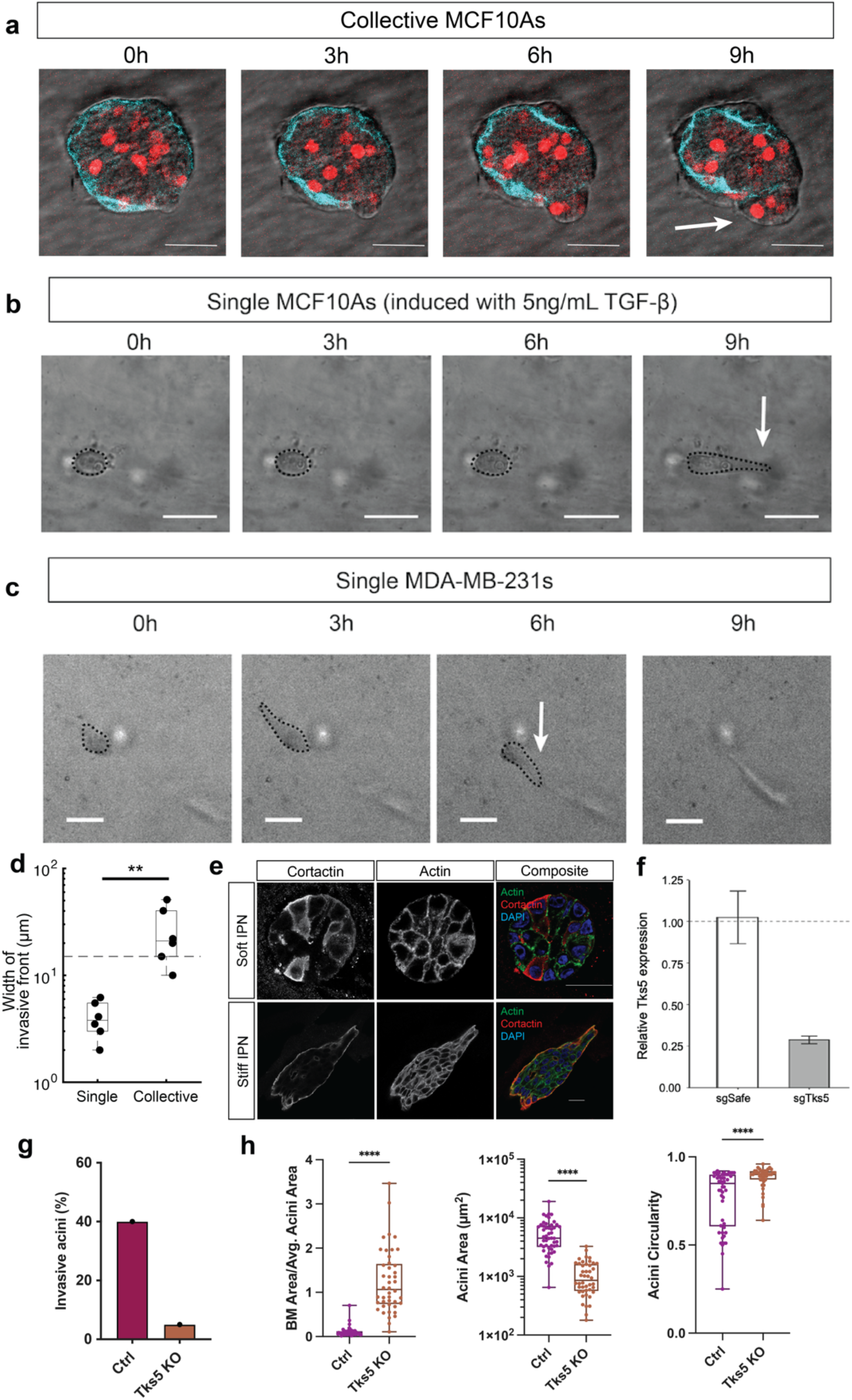
Single cells extend invadopodia like protrusions during invasions whereas collectively invading acini do not. **a,** Representative collective cluster of MCF10A migrating in stiff IPN. Scale bars, 50 μm. Arrow points to collectively migrating cell. Cyan = basement membrane, Red = cell nuclei. **b,** Representative single MCF10A induced with 5ng/mL TGF-β migrating in stiff IPN. Arrow points to protrusion. Scale bars, 50 μm. **c,** Representative single MDA-MB-231 migrating in stiff IPN. Arrow indicates direction of cell migration. Scale bars, 50 μm. Arrow points to protrusion. **d,** Width of invading front of single cells compared to collective cells \*\**p* < 0.01, Welch’s t-test. **e,** Cortactin (red), actin (green) and DAPI (blue) staining in acini of soft and stiff IPNs. Scale bar = 25 μm. **f,** RT-PCR of sgTks5 and sgSafe guides used in in CRISPR Cas9 MCF10As. **g,** Percentage of invasive acini in RFP-LifeAct MCF10As and CRISPR Tks5 KO MCF10As cells in one experiment. **h,** Acini area, acini circularity, and BM Area/Avg. Acini Area of RFP-LifeAct MCF10As and CRISPR Tks5 KO MCF10As cells in stiff IPNs. \*\*\*\**p* < 0.0001, Welch’s t-test.

### Global cell volume expansion mediates invasion

As invadopodia do not mediate collective invasion, we investigated other possible mechanisms by which cells invade collectively through the BM, starting with cell volume expansion as a potential mediator. Recent studies have demonstrated the involvement of cell volume changes in mechanotransduction and stem cell differentiation, and an increase in cell volume was found to be associated with invasion in a breast cancer 3D culture model that did not include an endogenous BM.^35–38^ Quantification of the cross-sectional area of acini and BM-enclosed area, expected to be correlated with volume, prior to BM breaching events showed an increase in both acini and BM area over time (Fig. 4a). After BM breaching, BM area decreases while acini area continues to increase (Fig. 4a). Inhibition of cell proliferation with mitomycin C decreased invasive events, although invasion still occurred (Fig. S5). These observations motivated analyses of cell volumes of invasive acini in stiff IPNs and the cell volumes of non-invasive acini in soft IPNs. Overall acini volumes in invading acini in stiff IPNs were greater than those in non-invading acini in soft IPNs (Fig. 4b, c). As overall acini volume changes can result from changes in cell number as well as cell volume, individual cell volumes were analyzed and found to be greater in the invading acini in stiff IPNs compared to non-invading acini in soft IPNs (Fig. 4d). A recent study investigating cell and nuclear volumes in invading MCF10A cell clusters in alginate-rBM IPNs, found that invasive cells exhibited greater cell and nuclei volumes compared to the core.^23^ In contrast, nuclear volume was similar across all conditions (Fig. S6) and local changes in cell volume in the invasive front compared to the non-invasive region of the cluster were not observed (Fig. S7). Instead, we observed increased cell volume globally in invasive acini compared to non-invasive acini. These findings demonstrate that cell volume expansion is associated with invasion.

**Figure 4.**
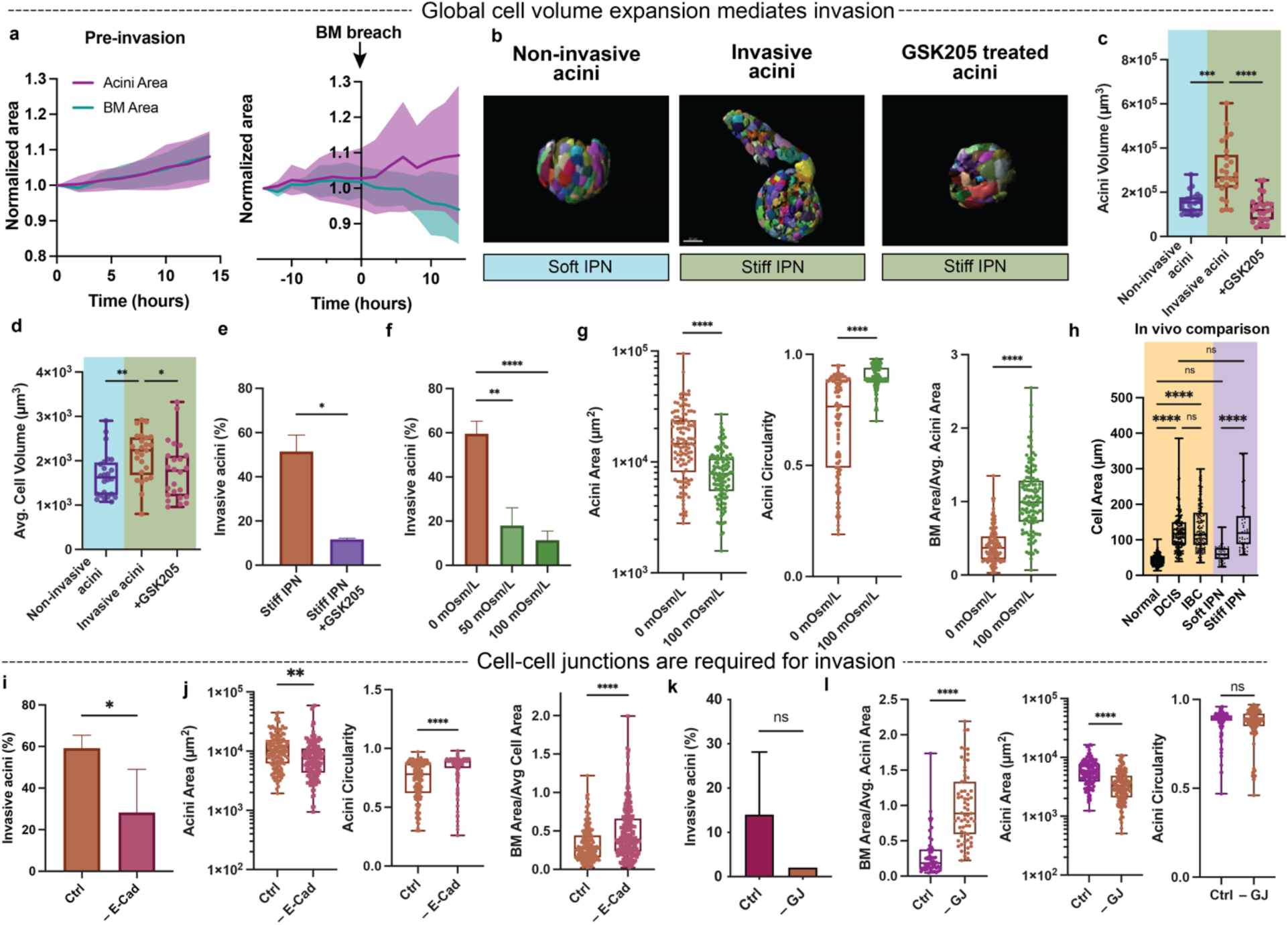
Global cell volume expansion and cell-cell junctions are required for invasion. **A,** Normalized area increases of acini (purple) and BM (blue) during pre-BM invasion (left) and BM breaching (right). **B,** Representative Imaris renderings of invasive acini, non-invasive acini, and GSK205 treated acini. Blue denotes soft IPN and green denotes stiff IPN. **C,** Acini volume of invasive acini, non-invasive acini, and GSK205 treated acini. \*\*\**p* = 0.0001, \*\*\*\**p <* 0.0001, Kruskal-Wallis with Dunn’s multiple comparisons test. **D,** Average cell volumes of invasive acini, non-invasive acini, and GSK205 treated acini. \**p* < 0.05, \*\**p* < 0.01, Kruskal-Wallis with Dunn’s multiple comparisons test. **E,** Percentage of invasive cells membrane in stiff IPNs and stiff IPN with GSK205, \**p* < 0.05, Welch’s-test. **F,** Percentage of invasive acini in stiff IPNs with varying osmotic pressure. ANOVA. \*\**p* < 0.01. **g,** Acini area, acini circularity, and BM area/avg. Acini area in stiff IPN with 0 and 100 mOsm/L osmotic pressure. \*\*\*\**p* < 0.0001, Welch’s t-test. **H,** Cell area based on E-cadherin stains in human breast tissue samples of normal, DCIS, and IBC conditions. \*\*\*\**p* < 0.0001, ns = not significant, one-way ANOVA with Tukey’s multiple comparisons test. Orange denotes in vivo and purple denotes in vitro. **i,** Percentage of invasive acini in stiff IPNs with E-cadherin function blocking antibody. \**p* < 0.05, Welch’s test. **j,** Acini area, acini circularity, and BM area/avg. Acini area in stiff IPN with control or E-cadherin function blocking antibody. \*\**p* < 0.01, \*\*\*\**p* < 0.0001, Welch’s t-test. **k,** Percentage of invasive acini in stiff IPNs with gap junction inhibitor. ns = not significant, Welch’s test. **l,** Acini area, acini circularity, and BM area/avg. Acini area in stiff IPN with control or E-cadherin function blocking antibody. ns = not significant, \*\*\*\**p* < 0.0001, Welch’s t-test.

Next, we tested the role of cell volume expansion in driving invasion. To pharmacologically perturb cell volume, transient receptor potential vanilloid-4 (TRPV4) ion channels were deactivated with GSK205, a small molecule antagonist. TRPV4 is a stretch activated ion channel that has been shown to mediate cell volume expansion through regulation of osmolality of calcium ions in the cytoplasm.^36, 39^ Volumes of invasive acini in stiff IPNs were greater than the volumes of acini in stiff IPNs treated with GSK205 (Fig. 4c). The average cell volumes within the invasive acini in stiff IPNs were also greater than the average cell volume in stiff IPNs treated with GSK205 (Fig. 4d). Treatment of acini in stiff IPNs with GSK205 decreased both the percentage of invasive cells and the percentage of breached BM (Fig. 4e). In addition to inhibiting cell volume expansion by blocking TRPV4 activity, cell volume expansion was also inhibited by applying osmotic pressure to the acini through the addition of 400 Da polyethylene glycol (PEG) into the cell culture media.^36^ Increasing PEG concentration in the media decreased the percentage of invasive acini (Fig. 4f). Collective cell invasion and BM breaching decreased with the addition of PEG as indicated by acini area, circularity, and ratio of BM to acini areas (Fig. 4g).

To establish the relevance of this putative volume expansion mechanism to invasive breast carcinoma, we analyzed the size of cells in samples from human patients. Cell areas in normal, breast carcinoma in situ (DCIS), and invasive breast cancer (IBC), showed that individual cell areas of breast ducts were smaller in normal tissue compared to DCIS and IBC (Fig. 4h). This matches with the finding that cell volumes are smaller in non-invasive acini compared to invasive acini. The cell areas of the breast ducts in normal tissue were similar to the cell areas of the acini in soft IPNs, and the cell areas of breast ducts in DCIS and IBC were similar to the cell areas of acini in stiff IPNs (Fig. 4h). This further demonstrates the similarity in cell morphology in soft and stiff IPNs to normal and cancerous breast tissue, respectively. Interestingly, the cell sizes between DCIS and IBC were similar, suggesting that cell volume expansion is necessary, but not sufficient to drive invasion on its own. Since cell volume was not significantly different between DCIS and IBC, we sought to examine other factors potentially promoting this invasive transition.

Experiments to perturb cell-cell adhesions demonstrated that cell collectivity is another factor that is necessary for BM and cell invasion. E-cadherin is a well-studied cell adhesion protein that physically link cells and facilitate transmission of signals between cells.^40^ Gap junctions enable the exchange of ions and fluid between cells and have been shown to regulate cell volume.^41^ Addition of a function blocking E-cadherin antibody reduced collective cell invasion and breaching of the BM (Fig. 4i, j). Addition of a gap junction inhibitor, carbenoxolone, also reduced BM breaching, though the reduction in the percentage of invasion acini compared to control was not significant (Fig. 4k, l). These pharmacological perturbations point to the importance of cell-cell adhesions in BM breaching and cell invasion.

### α3β1 integrin and actomyosin contractility drive invasion

Next, we explored the role of cell-generated contractile forces in BM breaching and invasion. Contractile forces from cells are generated by the actomyosin network and exerted on the surrounding matrix through integrin-based adhesions to matrix proteins. Since cells attach to the laminin in the BM via the α-chain of laminin, and as laminin densification and deformation was associated with invasion, we probed whether disrupting this link would prevent BM breaching and cell invasion.^3^ The addition of an anti-laminin α3 antibody to the culture media decreased the percentage of invasive acini and breached BM in stiff IPNs (Fig. 5a, b). α3β1 integrin has been shown to be recruited to focal contacts in cells, linking the BM to the actin cytoskeleton.^42^ Inhibition of α3 and β1 integrins separately and together significantly reduce the percentage of invasive acini (Fig. 5c, d). These data demonstrate that these specific binding sites of laminin α3 and α3β1 integrins are necessary for cell invasion and BM breaching.

**Figure 5.**
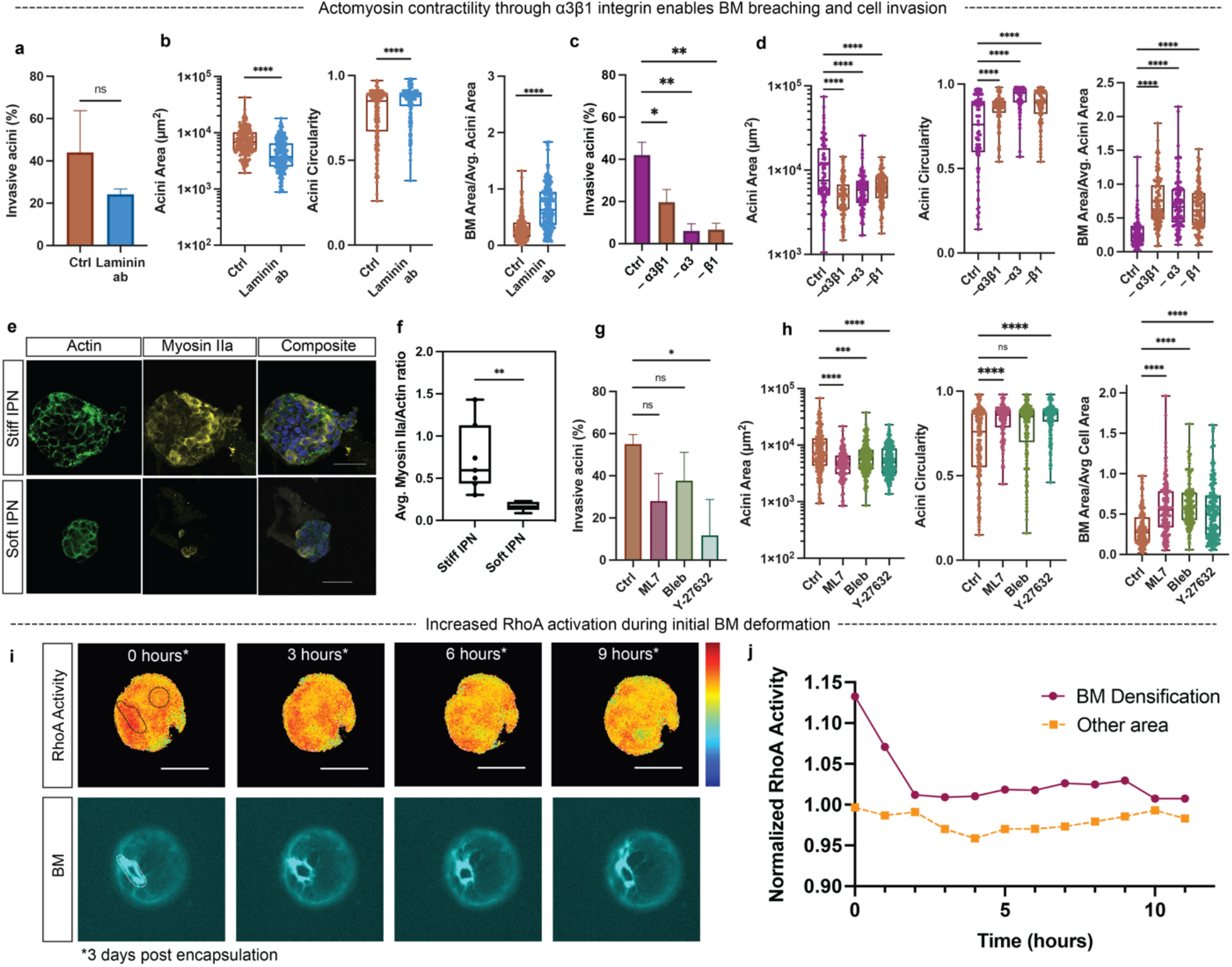
α3β1 integrin and actomyosin contractility enable BM breaching and collective invasion. **a,** Percentage of invasive acini in stiff IPN control or stiff IPN control with laminin antibody, ns = not significant, Welch’s-test. **b,** Acini area, acini circularity, and BM Area/Avg. Acini Area in stiff IPN control or stiff IPN with laminin antibody. \*\*\*\**p* < 0.0001, Welch’s t-test. **c,** Percentage of invasive acini in stiff IPN control, or α3 antibody, β1 antibody, or both α3 and β1 antibodies. \**p* < 0.05*, **p* < 0.01. Brown-Forsythe and Welch ANOVA tests. **d,** Acini area, acini circularity, and BM Area/Avg. Acini Area in stiff IPN control or stiff IPN with control, or α3 antibody, β1 antibody, or both α3 and β1 antibodies. \*\*\*\**p* < 0.0001, Welch’s t-test. \**p* < 0.05, \*\**p* < 0.01, Welch’s test. **e**, Actin (green) and myosin iIa (yellow) stains in soft and stiff IPNs. Composite images include actin, myosin iIa and nuclei (blue). Scale bar = 50 μm. **f,** Average Myosin iIa/Actin ratio in stiff and soft IPN. \*\**p* < 0.01, Welch’s t-test. **g,** Percentage of invasive acini in stiff IPNs with the addition of control, ML-7, blebbistatin, or Y-27632. \*\**p* < 0.01, ns = not significant. Brown-Forsythe and Welch ANOVA tests. **h,** Acini area, acini circularity and BM Area/Avg. cell area in stiff IPNs with the addition of control, ML-7, blebbistatin, or Y-27632. \*\*\*\**p* < 0.0001, \*\*\**p* < 0.001, ns = not significant. Brown-Forsythe and Welch ANOVA tests. **i,** RhoA FRET/donor ratio signal and BM signal over 9 hours after encapsulation. Scale bar = 50 μm. Warmer colors correspond to higher relative RhoA activity**. j** Normalized activity **of** RhoA at BM densification area (purple, solid) and other area (orange, dotted) divided by the average intensity of the entire cell area over 11 hours.

After finding the role of the integrin-laminin link on mediating invasion, we examined the role of actomyosin-based contractility. Quantification of the average myosin IIa/actin ratio pointed to increased staining of myosin IIa in stiff IPNs, suggesting increased acini contractility in stiff IPNs (Fig. 5e, f). Inhibition of myosin light chain kinase, myosin II, and Rho GTPase was conducted using ML-7, blebbistatin, and Y-27632, respectively. A reduction in cell invasion and BM invasion was found for all treatment conditions and by all metrics except for one (Fig. 4g, h). Next, the activation of RhoA was directly examined using a RhoA FRET biosensor to detect active RhoA (Fig. 5i, j).^43^ An initially high Rho FRET signal around the area of BM invasion was observed and followed by a sharp decrease after the BM hole opened up. The RhoA signal around the area of BM densification was higher than the RhoA signal around a non-breached area in the BM (Fig. 5j). This suggests that Rho activated contractility is involved in initial formation of a hole in the BM, followed by a relaxation back to the baseline. Together, these experiments illustrate the role of cell contractility in the BM breaching and collective cell invasion.

### Collectively invading cells exert tangential forces on BM

After elucidating the molecular mechanisms by which cells can breach the BM, we further explored the biophysical mechanism through which cells tear the BM to initiate invasion. Our data indicated that forces from increased acini volume and local cell contractility deform BM, which is followed by eventual tearing of the BM. To test the role of local cell contractility in deforming BM, local BM deformations in invasive acini were quantified using digital image correlation and both radial and tangential deformations in the BM pre-invasion were observed (Fig. 6a). When normalized by the total deformation magnitude, tangential deformations in the BM were larger than inward radial deformations or indentations (Fig. 6b, c), suggesting that local cell contractility dominantly deforms BM in the tangential direction. In agreement with this finding, BM tearing and hole dilation occurred primarily within the BM plane (Fig. 6d, Video 7). When investigating the post-BM breaching behavior of the cells, in which the cells invaded into the IPNs and BM indentation is observed, most of the difference in direction of cell invasion and laminin indentation were within 10 – 20° (Fig. 6d). This supports the finding that BM breaching enables cell invasion into the surrounding matrix. Observations of the cell movement trajectories immediately before BM indentation demonstrated an inward movement that corresponded to the subsequent negative BM curvature (Fig. 6f). Cells in acini with breached BM collectively exhibited greater radial velocities and decreased tangential velocities compared to cells in acini with intact BM (Fig. 6g, Fig. S8). Together, these findings highlight the movement of collective cells before and after breaching the BM, specifically demonstrating that cells exert tangential forces onto the BM prior to BM invasion.

**Figure 6.**
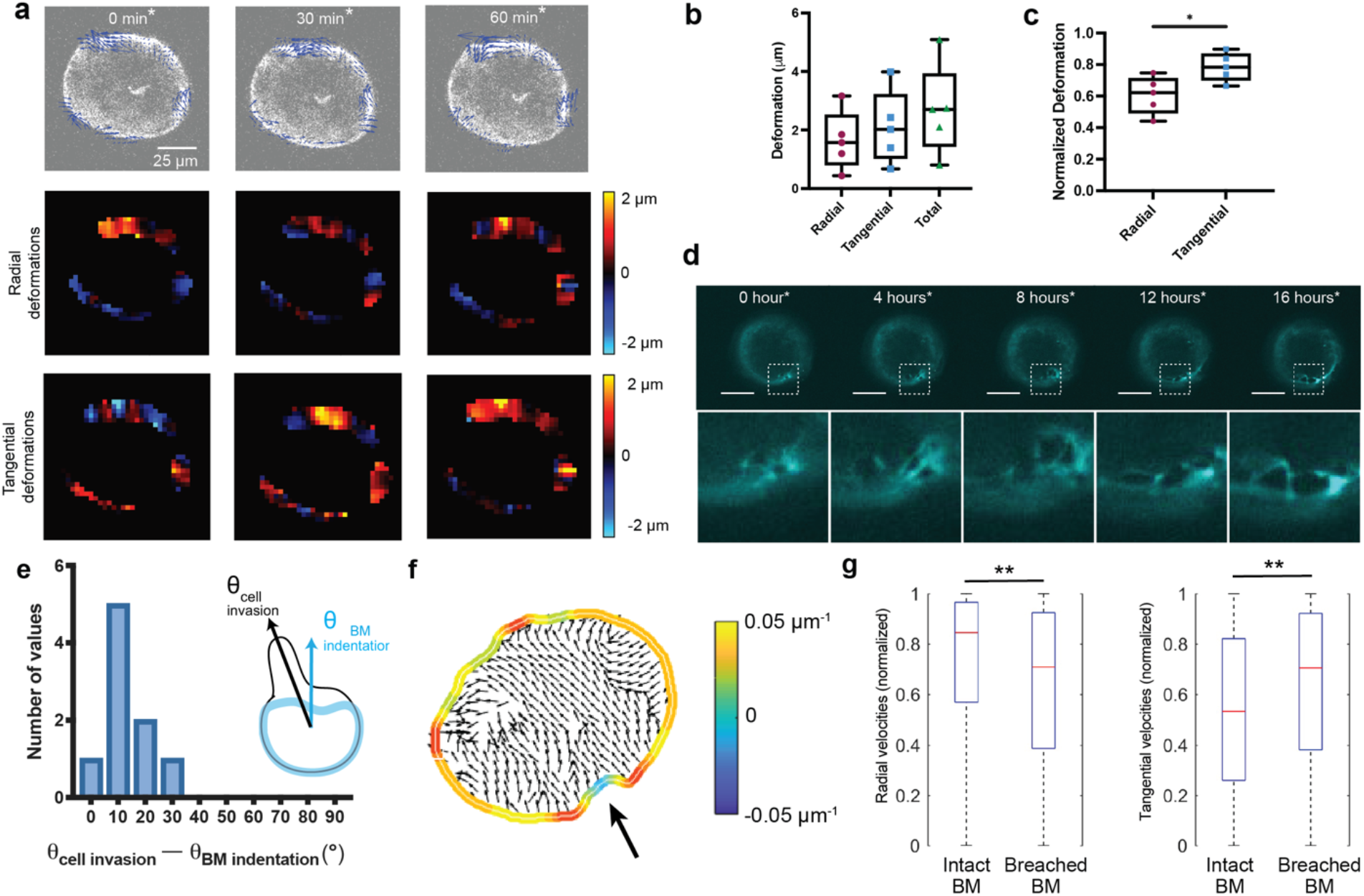
Collectively invading cells exert in-plane forces on the BM. **a,** Confocal images of BM at different time points before invasion (top). Heatmap of radial deformations of the BM at different time points before invasion (middle). Heatmap of tangential deformations of BM at different time points before invasion (bottom). **b,** Average of absolute radial and tangential deformations, and total deformation of BM. **c,** Normalized tangential and radial deformations with respect to total deformation of the BM. Each point corresponds to the average of absolute magnitudes over space and time for each acini. **d,** Representative time lapse images of BM deformation showing tangential hole opening in the boxed square, Scale bar = 50 μm. **e,** Histogram of differences between the angle between cell invasion and laminin indentation. **f,** Direction of cell movement within the acini compared to BM curvature deformation. Arrow in points to area of BM deformation. **g,** Tangential cell velocities and radial cell velocities, normalized by the total velocity, in acini with intact BM and breached BM. \*\**p* < 0.01, t-test.

### Global cell volume expansion and local tangential contractility breaches the BM

To examine how global cell volume expansion and tangential contractility could combine to drive BM breaching, we applied computational modeling. The model consisted of a growing tumor, surrounded by a thin BM and IPN matrix (Fig. 7a, b). Cells at the BM interface apply tangential contractile forces to the BM at either a baseline, resting level or in an activated state (Fig. 7a, b). Global cell volume expansion drives pre-stretching of the BM with higher tangential stresses on the BM relative to radial stresses (Fig. 7c). Local cell contractility alone, in the absence of global volume expansion, leads to high tangential BM stresses in the region of the activated cell. Finally, when global cell volume expansion is combined with activated local cell contractility, the highest tangential stresses in the BM are observed, which could potentially lead to breaching of the BM. While there is uncertainty in some of the parameters, the model illustrates how expansion combines with contractility to lead to high tangential stresses on the BM. Interestingly, the maximum stress in the case of volume expansion and local cell contractility combined exceeds the sum of the maximum stresses in the case of each behavior on its own, highlighting the synergy generated between these two mechanisms. Together the modeling and experiments indicate that cell volume expansion acts synergistically with activated local cell contractility to drive rupture of the BM and facilitate invasion.

**Figure 7.**
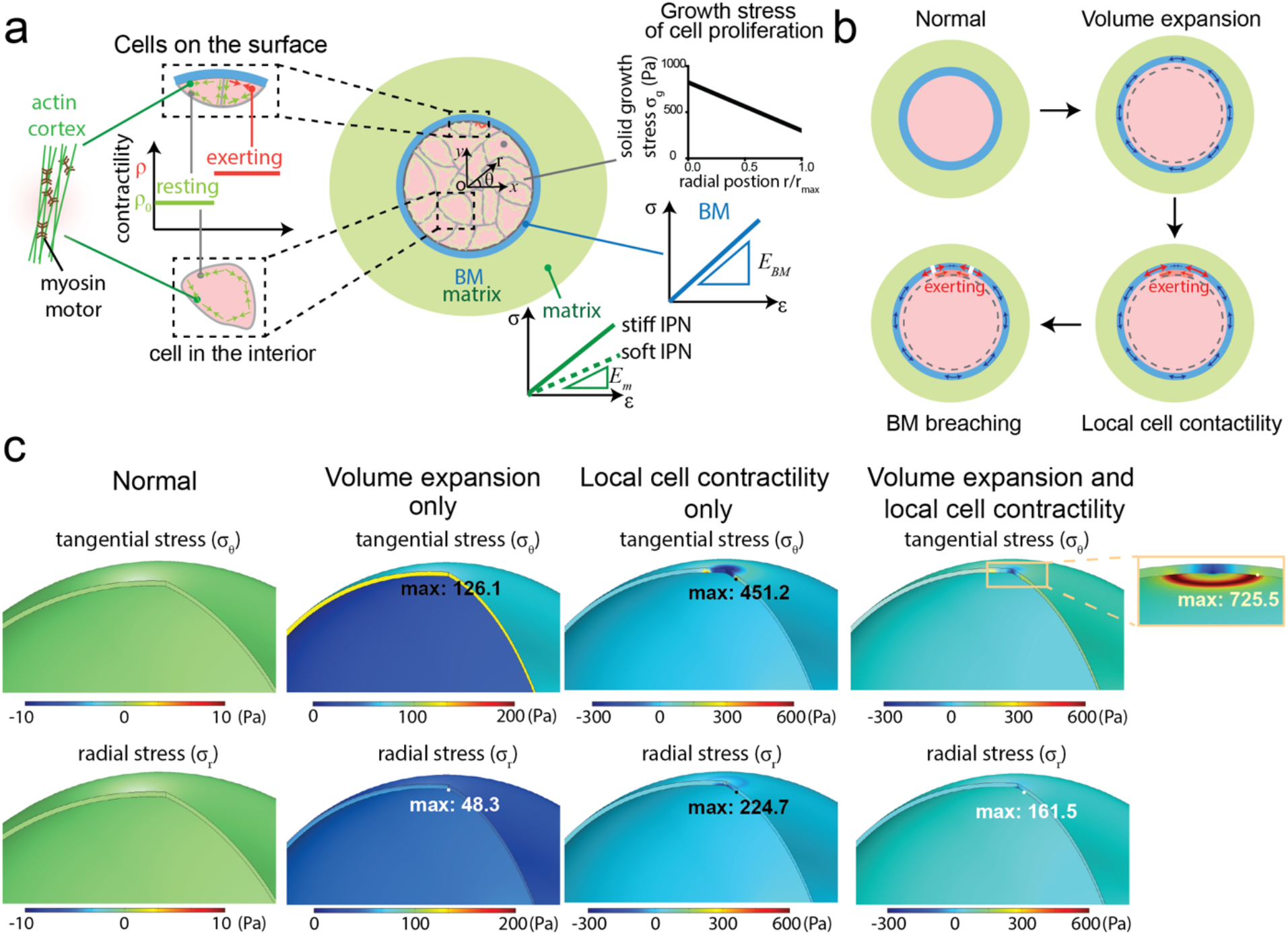
Global cell volume expansion and local tangential contractility combine to drive BM breaching. **a**, A schematic highlighting the process of force-driven BM breaching. Cell proliferation and cell volume expansion induces the growth stress and overall volume expansion in the tumor (black curve). Cells typically exhibit baseline levels actomyosin contractility on the BM (black curve), but, at increased stiffness, some cells exhibit enhanced actomyosin contractility. The IPN hydrogels and BM are considered to be linear elastic for this model (green and blue curves). **b**, A schematic of BM rupture due to tumor expansion and enhanced local contractile forces: normal tumor, volume expansion inducing tangential stress in the BM, local cell contractility resulting in higher tangential stress in the BM, and BM breaching due to higher tangential stress. **c**, computational modeling indicates BM stresses tangential to the BM and in the radial direction that are expected to occur due to volume expansion only, local cell contractility only, and the combination of volume expansion and local cell contractility (as indicated by heatmaps). Maximum stresses for each case are written in the figure.

## Discussion

In this study, we developed a 3D *in vitro* model of breast carcinoma invasion to elucidate how protease-dependent and protease-independent modes of invasion contribute to collective invasion of the endogenous BM. We find that both protease and force driven mechanisms were important in the BM breaching process (Fig. 8). BM invasion was driven by global increase in cell volumes and local cell contractility and tangential forces on the BM, which were mediated by laminin α3 and α3β1 integrin interactions between the cell and BM.

**Figure 8.**
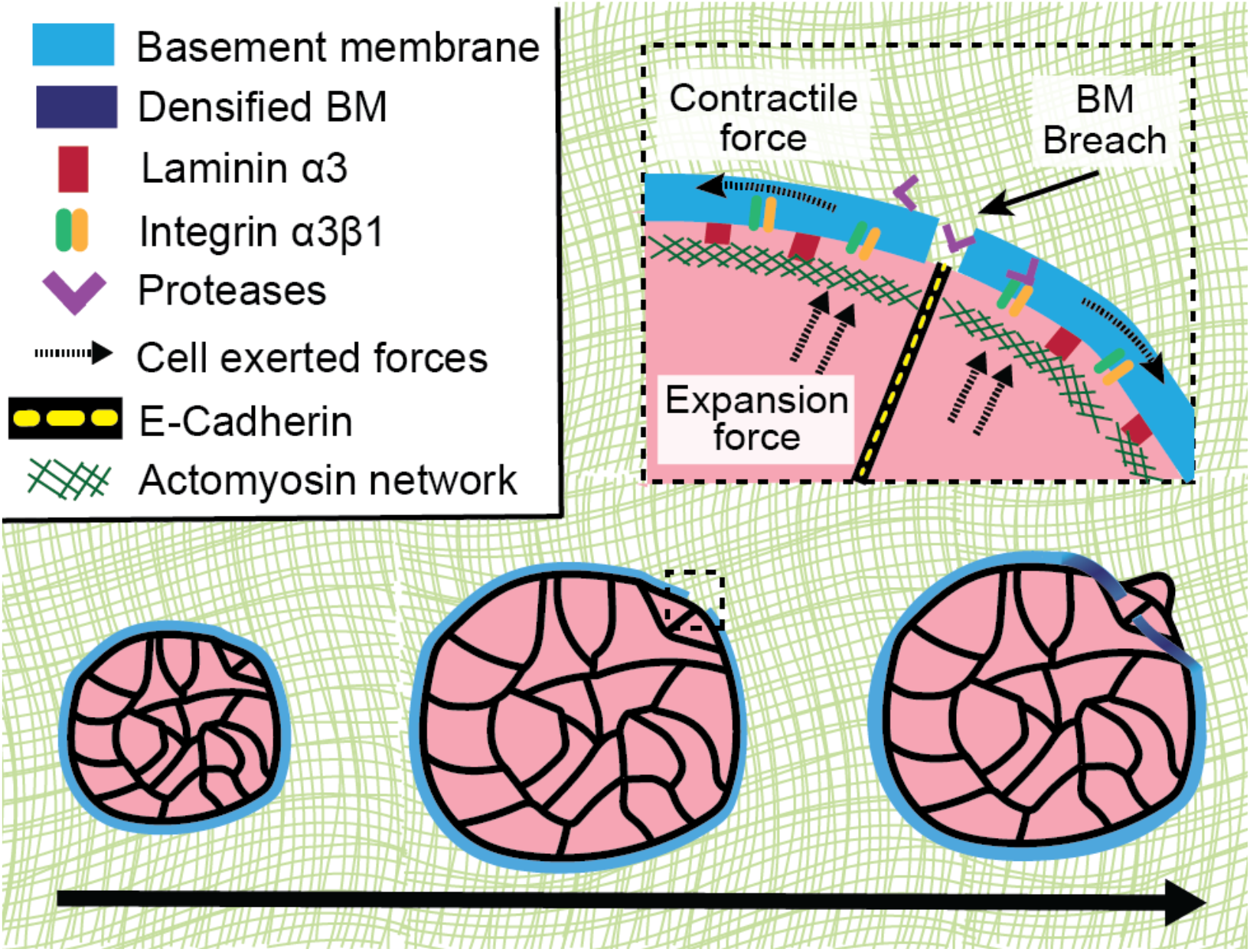
Schematic of basement membrane invasion process by mammary epithelial cells in a 3D *in vitro* model of breast carcinoma. Schematic highlighting the process of protease-dependent and force-driven BM Invasion by collective mammary epithelial cells. Protease activity combined with both cell volume expansion and tangential forces exerted by cells locally on the BM through actomyosin contractility enable breaching through the BM. Laminin α3 and α3β1 integrins drive the cell-BM attachments to physically pull in on the BM, leading to densified areas of BM around the cell invasion site.

Our 3D culture model recapitulated key features of the physiological BM and breast carcinoma invasion. Separating the primary tumor from the surrounding stromal matrix, the BM is rich of mechanical cues. The two primary components of BM are laminin, which enables cell adhesion to the BM, and collagen-IV, which provides structural integrity to the BM.^2, 3^ While the BM is nanometers to micrometers thick, it is nanoporous and measurements indicate the elastic modulus (i.e. “stiffness”) of BM to be in the kPa to mPa range, though also exhibiting nonlinear elasticity and strain stiffening.^44–48^ However, the accuracy of BM stiffness measurements is uncertain due to the challenges of measuring stiffness of such thin materials and potential variability in BM mechanics between tissues and animals. BM thickness in our *in vitro* model was similar to BM thickness in DCIS breast tissue samples, while BM thickness in normal breast tissue samples was significantly greater. Though we are unable to examine the molecular crosslinking and architecture of the BM, the similarity of BM thickness and localization of laminin and collagen-IV compared to *in vivo* samples supports the relevance of our model to the DCIS to invasive breast cancer transition. Computational and *in vitro* experiments have shown that cells can sense stiffness on the micron scale, signifying that cells can not only sense the mechanical properties of the immediate BM, but they can also sense the mechanical properties of the surrounding stromal matrix.^28, 49^ Thus, acini became invasive in stiff IPNs and remained non-invasive in soft IPNs. Cells are likely able to sense the IPN stiffness through the strong mechanical connectivity between endogenous BM and the IPN mediated by binding interactions between collagen-IV on the outer layer of endogenous BM with laminin and collagen-IV in the rBM in the IPN.

Our study was in part motivated by previous studies demonstrating that force is important for BM invasion, and our results are consistent with this notion, challenging earlier perspectives that the BM invasion is exclusively driven by protease degradation via matrix metalloproteinases (MMPs).^50^ Early studies revealed that cancer cells can degrade the collagen-IV in BM and greater degradation correlated with greater metastatic potential.^9^ Several studies have since identified specific MMPs that facilitate tumor cell invasion through the BM.^10, 51^ More recent studies have revealed that cells can use protease-independent, force driven mechanisms to invade through the BM. Individual cancer cells can use invadopodia to exert force to open up holes in nanoporous BM-like matrix to enable invasion, independent of proteolytic degradation, if the matrix exhibits sufficient matrix mechanical plasticity.^12^ Individual anchor cells in *Caenorhabditis elegans* can use force to invade through the BM.^13, 52^ Further, cancer-associated fibroblasts pull, stretch and soften the BM to create holes that cancer cells can invade through, although direct *in vivo* evidence of this mechanism is lacking.^14^ Our study demonstrated that both protease and force are important in mediating BM invasion. While protease inhibition decreased invasion, it did not completely abrogate invasion, indicating the presence of protease independent modes of invasion. Accumulation of laminin signal around the BM invasion sites also points towards the role of forces versus proteases since proteases would be expected to diminish the BM signal, as opposed to increasing it. Moreover, we observed increased actin and pFAK, two key mediators of force generation, around the area of BM densification. These openings in the BM barrier persist and do not close, demonstrated mechanical plasticity of the BM, as demonstrated by experiments using latrunculin to depolymerize actin. Together, these results show that cells use a combination of proteases and forces to breach the mechanically plastic BM barrier.

Cells collectively invade during the initial breaching of the BM and the inherent mechanics of collective migration in this context differ substantially from collective migration in type-1 collagen rich matrices or single cell movements. Cell structures such as filopodia and invadopodia have been implicated in single cell migration, but were not observed here.^12, 53^ Cell-cell adhesions within collective cells give rise to synergistic interactions and migratory behaviors that differ from single cell migration.^54, 55^ While studies of cells migrating as collective monolayers offered novel insights into cell migration mechanics, the 2D environment does not require cells to interact and overcome a physical barrier, a critical feature of BM invasion.^56, 57^ A study investigating 3D matrix confinement in collagen-I, the primary component of stromal matrix, showed multiple collective streams of cells migrating outward from the cell spheroid and single cell detachment occurring at low collagen-I densities.^21^ At higher collagen-I densities, cells only migrated collectively, although significantly fewer collective streams of cell migration were detected.^21^ In contrast, we typically observe one point of collective invasion from the acini in the IPNs and typically do not observe single cell detachment. This is likely due to the nanoporosity of the IPNs, which is more confining compared to microporous collagen-I gels. Another study in collagen-I gels observed leader cell behavior during collective migration in collagen-I gel, as indicated by increased cytokeratin-14 (K14) and p63 expression by the leading cells.^22^ In our system, normal acini in soft IPNs exhibited heterogenous expression of K14 and invasive acini in stiff IPNs exhibited overall greater K14 expression, but not specifically at the invasive front (Fig. S9). These seemingly disparate results suggest the possibility that leader cell behavior is not established until cells are collectively migrating within the collagen-I-rich stromal matrix, following BM invasion. In summary, our 3D *in vitro* model of breast carcinoma invasion demonstrated unique collective migration behavior, involving collective invasion of the BM at a single focal point and without a K14-presenting leader cell.

Here, we gleaned a new mechanism of collective cell migration that involves the use of global cell swelling combined with local contractility to rupture the endogenous BM to facilitate invasion. The use of cell swelling to breach the BM is a collective phenomenon, since a single cell swelling by itself would likely not be enough to rupture the BM. This points to the idea that collective cells utilize mechanisms unique from those of single cells for generating force and breaching the BM, due to increased ability to generate greater forces and/or deformations. We show that α3β1 integrins, which play a role in focal contact adhesions to the BM, are necessary for BM invasion and cell invasion. Other integrins, including α6β4 integrins, which are found in hemidesmosomes that attach to the BM in normal mammary epithelium, connecting the BM to the keratin intermediate filament network, might also play a role though they were not specifically investigated here. We expect that α6β4 integrins might play a primary role in mechanotransduction, or stiffness sensing, but perhaps not the physical process of invasion.^25, 58^ α3β1 integrins are likely to play a key role in transmitting forces from cell contractility to the BM, though they could also play a role in connecting cell swelling to the BM as well. It is possible that other cells, such as cancer associated fibroblasts,^14^ in the tumor microenvironment contribute to the weakening of the BM through contractile means. Additionally, interfacing luminal epithelial cells and the BM are myoepithelial cells, which exhibit contractile abilities.^59^ Even beyond primary tumor escape, collective cells may retain this survival advantage throughout the rest of the metastatic cascade, implicating collective dissemination as an important feature of metastasis.^60^ Our results seemingly contrast past studies that showed that E-cadherin inhibition leads to increased metastasis. However, since different stages of metastasis likely require distinct migration strategies, it is possible that advantage conferred by E-cadherin changes during the various stages of metastasis and becomes less important in the later stages of invasion.

In conclusion, this study reveals that both proteases and forces generated by cells are critical for BM invasion. This suggests that therapeutics only targeting proteases may not be enough to inhibit initial BM invasion; alternative therapeutic approaches could include inhibiting cellular contractility, limiting volume expansion, and modulating the mechanical properties of the BM to prevent cancer cell invasion. More generally, these insights into initial invasion in breast cancer may extend to other carcinomas or diseases characterized by impaired BM function and in developmental biology.

## Methods

### Cell culture

MCF10A human mammary epithelial cells (ATCC) were cultured in Dulbecco’s modified Eagle’s medium/Nutrient Mixture F-12 (DMEM/F12) medium (Thermo Fisher) supplemented with 5% horse serum (Thermo Fisher), 20 ng/ml EGF (Peprotech, Inc.), 0.5 µg/ml hydrocortisone (Sigma), 100 ng/ml cholera toxin (Sigma), 10 µg/ml insulin (Sigma), and 1% penicillin/streptomycin (Thermo Fisher). MCF10A cells expressing RFP-LifeAct were used for live-imaging experiments. MDA-MB-231 human breast cancer adenocarcinoma cells (ATCC) were cultured in high-glucose DMEM (Thermo Fisher) supplemented with 10% fetal bovine serum (Hyclone) and 1% penicillin/streptomycin. Cells were routinely split every 3-4 days with 0.05% trypsin/EDTA and cultured in a standard humidified incubator at 37°C and 5% CO_2_.

### Tissue immunohistochemistry

Immunofluorescence (IF) staining was performed on paraffin-embedded tissue microarray (TMA) sections (Stanford TA 417, 419, and 424). Primary antibodies used for IHC staining include Laminin-332 (Santa Cruz Biotech Cat# sc-28330) and E-cadherin (BD Pharmingen Cat# 560062 conjugated with af647). TMA sections measuring 4 µm were deparaffinized in 3 changes of xylene for 10 mins each and hydrated in gradient series of ethyl alcohol. For E-Cadherin, target retrieval in 10 mM citrate pH6 (Agilent, Santa Clara, CA, catalog #S2369) was performed to retrieve antigenic sites at 116 °C for 3 min. For Laminin-332, retrieval was done with Proteinase K (Agilent, Santa Clara, CA, catalog #S3020) for 15 min at room temperature and visualized with anti-mouse IgG1k af647 (Invitrogen/Life Technologies Eugene, OR catalog #A21240). All samples were counterstained with DAPI (Invitrogen/Life Technologies, Eugene, OR catalog #P36935)

### Alginate preparation

High molecular weight sodium alginate rich in guluronic acid blocks (FMC Biopolymer, Protanal LF 20/40, High-MW, 280 kDa) was dialyzed against deionized water for 3 – 4 days using dialysis tubing with a molecular weight cutoff of 3.5 kDa. Alginate was subsequently treated with activated charcoal, sterile filtered and lyophilized. Prior to experiments, lyophilized alginate was reconstituted to 2.5 wt% in serum-free DMEM/F12 (Thermo Fisher).

### Acini formation

6-well plates were coated with 150uL of rBM (Corning) in 6-well plates were placed into 37°C incubator with CO_2_ to gel for at least 10 minutes. To form MCF10A acini in the 6-well plates, MCF10A cells were trypsinized, strained through a 40 µm cell strainer to enrich for single cells, counted on a Vi-CELL (Beckman Coulter Life Sciences), and seeded at a final concentration of 1 × 10^5^ cells/ml hydrogel. Cells were cultured in 2 mL of complete DMEM/F12 medium supplemented with 2% rBM. Addition of 2% rBM is necessary to drive the development from single cells to acini.^24^ rBM supplemented complete media was replenished on day 4. Once the acini were formed, the acini were extracted by adding 2mL of ice-cold 50mM EDTA in PBS per well and scrapping off the layers with a cell scraper. After a 20-minute incubation on ice, the acini-containing EDTA mixture was spun down for 5 minutes at 500*g* at 4°C. The acini were resuspended in MCF10A resuspension media and centrifuged at 500*g* for another 5 min. After centrifugation, the supernatant was aspirated, and the acini was resuspended in DMEM/F12 media.

For live imaging experiments of BM, except FRET, Alexa Fluor 488 conjugated anti-laminin-5 antibody, clone D4B5 (Millipore Sigma) was added to the resuspended media at a 1:100 concentration and incubated on ice for one hour. For pharmacological inhibition experiments, inhibitors were added to the resuspended media and incubated on ice for one hour. After incubation, the cells were spun down for 5 minutes at 500*g* at 4°C, the supernatant was aspirated, and the acini was resuspended in DMEM/F12 media.

### Encapsulation of acini in hydrogels

IPNs were formed as previously described.^25^ 24-well plates were coated with 50μL of rBM (Corning) and allowed to gel for at least 10 min in a 37°C incubator with CO_2_. Standard 24-well plates (Corning) were used for immunohistochemistry and 24-well glass bottom plates (Cellvis) were used experiments that required sample imaging. Alginate was mixed with rBM, acini, and DMEM/F12 in a 1.5ml Eppendorf tube and added to a 1 ml Luer lock syringe (Cole-Parmer). Appropriate volume of each component were added to reach a concentration of 5 mg ml^-1^ alginate and 4.4 mg ml^-1^ rBM in the final IPNs. For experiments requiring fiducial marker beads, 0.2 μm dark red microspheres (Thermo Fisher) were added to syringe containing alginate at a 100-fold dilution. The second 1ml syringe contained DMEM/F12 containing 5mM CaSO_4_ for soft IPNs or DMEM/F12 containing 20mM CaSO_4_ for stiff IPNs. The two syringes were coupled with a female-female Luer lock (Cole-Parmer), quickly mixed, and instantly deposited into the 24-well plate coated with rBM. A transwell insert (Millipore) was immediately placed on top of the IPN to prevent floating and the IPNs containing acini were placed in a 37°C incubator with CO_2_ to gel for at least 30 minutes. After the IPNs have gelled, 1.5ml of complete media, with pharmacological inhibitors if required, was added.

### Acini and cell volume measurements

Imaris Bitplane was used to create surface renderings of RFP-LifeAct-transfected MCF10A cells. Images acquired during confocal imaging were imported into Imaris (Bitplane), a software that enables 3D volume rendering and analysis. Cell volume measurements were made using the RFP channel of the cells. For cell volume, the Cells program was used with the following parameters: cell smallest diameter = 10.00 μm, cell membrane detail = 2.00 μm, cell filter type = local contrast. For cell nuclei, the Surfaces program was used with the following parameters: surface grain size = 0.800 μm, and diameter of largest sphere = 5.00μm. Images were all manually filtered for abnormal cell or nuclei volumes.

### Mechanical Testing

Material stiffness and plasticity measurements were performed using an AR2000EX stress-controlled rheometer (TA Instruments, New Castle, DE) with 25-mm top- and bottom-plate stainless steel geometries. IPN solutions without cells were deposited onto the center of the bottom-plate. A 25 mm flat-top plate was quickly lowered to form a 25 mm disk before gelation of the IPN and mineral oil (Sigma-Aldrich, St. Louis, MO) was applied to the edge of the IPN to prevent sample dehydration. A time sweep was conducted at 37°C, 1 rad s^-1^ and 1% strain for 2 h and measurements were taken once the storage modulus reached equilibrium. The Young’s modulus (E) was calculated using the equation:

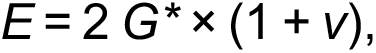

where the Poisson’s ratio, *v,* is assumed to be 0.5 and G* is the bulk modulus calculated from the storage (G’) and loss moduli (G’’) measured using:

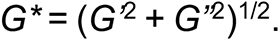

To characterize plasticity, a creep recovery test was performed immediately after the time sweep. Strain measurements were recorded as the IPNs were subjected to a constant shear stress (25 Pa) for 1 h, followed by a recovery period wherein the sample was unloaded (0 Pa). The resulting strain was recorded as a function of time after 10,000 s.

### Immunofluorescence for fixed cells

To fix hydrogel samples for immunofluorescence staining, hydrogels containing wild-type MCF10A cells were fixed for 30 min in warmed 4% paraformaldehyde in serum-free DMEM/F12 at room temperature. Gels containing acini were washed twice in PBS for 15 min, incubated in 30% sucrose solution in PBS overnight, and incubated in a 1:1 mixture of 30% sucrose and optimal cutting temperature (OCT) compound (Tissue-Tek) for at least 4 h. Gels were embedded in OCT and frozen on dry ice. Frozen gels were sectioned with a cryostat into ∼50 µm sections and adhered onto microscope slides. Slides were blocked in a solution of 10% goat serum (Thermo Fisher), 1% bovine serum albumin (Sigma), 0.1% Triton X-100 (Sigma) and 0.3 M glycine (Sigma) for 1 h at room temperature. Primary antibodies were diluted in the blocking solution (1:100) and incubated at 4 °C overnight. Alexa Fluor 488–phalloidin (1:100 dilution, Thermo Fisher) and 4′,6-diamidino-2-phenylindole (DAPI; 1 µg ml^−1^) were diluted in the blocking solution and incubated for 1 h. Fluorescently conjugated secondary antibodies were incubated in blocking solution for 1 hour at room temperature. The slides were then washed three times for 5 min in DPBS, and coverslips were applied with Prolong Gold antifade reagent (Thermo Fisher). Slides were imaged on a Leica SP8 laser scanning confocal microscope with a 63X NA1.40 PlanApo oil immersion objective. Antibodies used were anti-laminin-5 (γ2 chain, clone D4B5, Millipore Sigma, #MAB19562), anti-collagen IV antibody (Sigma, #SAB4500369), anti-non-muscle Myosin IIa antibody (Abcam, #ab55456), anti-phospho-FAK antibody (Tyr 397, Invitrogen, #700255), anti-E-cadherin (BD Biosciences, #610181), anti-β4 integrin (Thermo Fisher), Alexa Fluor 488 goat anti-mouse IgG_2a_ (#A21131), Alex Fluor 488 goat anti-mouse IgG2b (#A21141), Alexa Fluor 647 goat anti-mouse IgG_1_ (#A21240) and Alexa Fluor 647 goat anti-rabbit (#A21244).

For collagen-IV and laminin-332 staining endogenous BM, the acini were not embedded in a hydrogel since collagen-IV is also present in rBM. Instead, seven-day acini were seeded onto poly-lysine coverslips, fixed by adding 1mL formaldehyde for 20 minutes at room temperature, and washed twice with PBS. The same staining protocol as previously described was followed.

### Pharmacological inhibition

Pharmacological inhibitors were added to the cell media after the hydrogels were formed. Images were acquired three days later. The concentrations of pharmacological inhibitors used were: 50 μM Blebbistatin (Abcam, myosin II inhibitor), 10 μM FAK inhibitor (Selleck Chem), 25 μM GM6001 (Millpore, MMP inhibitor), 30 μM GSK205 (Calbiochem, TRPV4 antagonist) 2.5 μM Latrunculin A (Tocris Bioscience, actin polymerization inhibitor), 100 μM Marimastat (Tocris, MMP inhibitor), 25 μM ML-7 (Tocris, myosin light-chain kinase inhibitor), 500 μM carbenoxolone (Sigma, gap junction inhibitor) and 25 μM Y-27632 (Sigma, ROCK inhibitor). The protease inhibitor cocktail contained 20 μM Marimastat, 20 μM Pepstatin A (Sigma), 20 μM E-64 (Sigma), 0.7 μM Aprotinin (Sigma), and 2 μM Leupeptin (Sigma) as previously described.^30^ The concentrations of function blocking antibodies used were: 1 μg mL^−1^ Anti-laminin 5 antibody (R&D systems, #MAB2144), 1 μg mL^−1^ α3 anti-integrin antibody, clone P1B5 (Sigma, #MAB1952Z), 1 μg mL^−1^ β1 anti-integrin antibody, clone P5D2 (Abcam, #ab24693), 1 μg mL^−1^ α6 anti-integrin antibody, clone GoH3 (Sigma, #MAB1378), 1 μg mL^−1^ β4 anti-integrin antibody, clone ASC-8 (Sigma, #MAB2059) and 1 μg mL^−1^ E-cadherin antibody, clone DECMA-1 (Sigma, #MABT26). Appropriate controls used include IgG from mouse serum (Sigma, #I5381) and IgG from rat serum (Sigma, #I4131).

### Confocal microscopy

Microscope imaging was performed with a laser scanning Leica SP8 confocal microscope or a Nikon Ti2-E inverted microscope, both fitted with a temperature and incubator control suitable for live imaging (37°C, 5% CO_2_). A 10× NA 0.45 dry objective or 20× NA 0.75 dry objective was used for the Nikon microscope. A 10× NA 0.4 dry objective was used for the Leica microscope. For live-cell time-lapse imaging, images were acquired 3 days after acini were encapsulated in IPNs; images were acquired overnight every 30, 60, or 120 minutes. For pharmacological inhibition experiments, images were acquired 3 days after acini were encapsulated in IPNs.

### Basement membrane breaching and cell invasion quantification

Invasive acini and their BM structures were manually counted in at least 10 fields of views with at least 40 total acini counted in 3 experiments. Image stacks of 100 μm and 10 μm stack sizes were acquired. Invasive acini were identified as those in which collective cells were identified to have migrate from the acini in a 2D field of view. The maximum intensity projection was taken for each 100 μm image stacks and a Gaussian blur image filter was applied using ImageJ. Next, a threshold was applied to the images to convert the image from grayscale into a binary image. Further quantification of BM breaching and cell invasion parameters were also performed, including: a) acini area, b) acini circularity, and c) BM area/average acini area. The combination of a large acini area and low acini circularity are characteristic of invading acini. A BM area/average acini area close to 1 is characteristic of an intact BM that has not been breached, while a BM area/average acini area less than 1 is characteristic of a BM that has been breached.

### Osmotic pressure studies

To modulate osmotic pressure in the culture media, 400 Da polyethylene glycol (PEG 400, TCI America) was added as previously described.^36^ PEG 400 concentrations of 0% wt/vol (0 mOsm/L), 2% wt/vol (50 mOsm/L), and 4% wt/vol (100 mOsm/L) was added to the media after hydrogel formation.

### Förster Resonance Energy Transfer (FRET) experiments

Optimized biosensor for RhoA was developed based on a previously reported single-chain, genetically encoded FRET biosensor design (pubmed.ncbi.nlm.nih.gov/16547516/). Briefly, the original, dimeric ECFP and dimeric Citrine YFP fluorescent proteins were exchanged for the monomeric versions containing the A206K mutation (pubmed.ncbi.nlm.nih.gov/22869113/). The mECFP was further modified using synonymous codons to prevent random homologous recombination during stable genomic integration and expression in target cells (pubmed.ncbi.nlm.nih.gov/25877922/). The biosensor was further optimized to improve the dynamic range of FRET change by utilizing a circularly permutated (cp)-173 mECFP (Fig. S10a). The mcp173ECFP was produced by a 2-step PCR reaction using the following primer sets: 5’-GCTAATGCATATACAAGGATCCGGCATGGATGGAAGCGTGCAGCTGGCTGATC-3’, 5’-GGTTAATACATGTTAGAAGCTTTCAATATTATGCCTAATTTTAAAG-3’, 5’- TTGGACACCATGCCGCCGCTGCCGCCTTTATACAGTTCATCCATTCCCA-3’, and 5’-ACTGTATAAAGGCGGCAGCGGCGGCATGGTGTCCAAAGGAGAAGAACTGTT-3’. The combination of the monomeric mutations and the circular permutations resulted in approximately 4.5 fold improvement in the fluorometric response in FRET/donor ratio comparing the constitutively activated mutant Q63L (“on”) versus the dominant negative T19N (“off”) versions in live suspensions of HEK293T cells overexpressing the biosensor and measured in a spectrofluorometer (Fig. S10b). The improvement in the dynamic range was achieved both by an increased on-state FRET due to the optimized dipole coupling from the use of a circularly permutated mECFP, and an improvement in the fall-off of FRET during the off-state from the monomerization of the fluorescent proteins (Fig. S10c). The biosensor responded appropriately to various mutations as demonstrated previously for the original version of this biosensor (pubmed.ncbi.nlm.nih.gov/16547516/), as well as interaction with an upstream regulator Rho-guanine nucleotide dissociation inhibitor (GDI) (Fig. S10d). Transient expression of the constitutively activated Q61L versus the dominant negative T19N versions of the biosensor in MCF10A cells showed approximately 40% difference in the average whole-cell FRET/donor ratio in microscopy, while the relative ratio differences as measured within single cells indicated a local dynamic range of 1.0 to 3.4 in the pseudocolor scaling (Fig. S10e). The biosensor as designed does not compete against binding to other cellular targets of activated RhoA, which minimizes the overexpression artefacts in living cells (Fig. S10f).

The parental MCF10A cells were stably transduced with tetracycline-OFF-transactivator system (tet-OFF; Clontech) following a previously described protocol (pubmed.ncbi.nlm.nih.gov/27815877/). The biosensor expression cassette was transduced into the stable tet-OFF MCF10A cell line under the regulation of a tet-inducible minimal CMV promoter (pRetro-X-Puro; Clontech) and selected using puromycin for a stably integrated population. The biosensor expression was subsequently induced by withholding doxycycline for 48 hrs, and the cells were FACS sorted to enrich for the population of positive transductants. Biosensor expression levels upon induction in the FACS sorted population were analyzed by western blot, showing approximately 38% of the levels of the endogenous RhoA GTPase (Fig. S11a, b), similar to previously reported systems of this type (pubmed.ncbi.nlm.nih.gov/16547516/; pubmed.ncbi.nlm.nih.gov/21474314/). Induction of the biosensor expression in MCF10A cells did not affect the relative expression levels of endogenous Rho GTPases, including RhoA, Rac1 and Cdc42 (Fig. S11c-h).

MCF10A cells stably harboring the inducible RhoA biosensor were normally maintained in MCF10A growth media with 1ug/ml doxycycline, 1mg/ml G418, and 10 ug/ml puromycin. 48 hours prior to encapsulation of cells in IPNs, MCF10A with RhoA biosensors were trypsinized and plated in tissue culture plates without doxycycline. This process was repeated 24 hours prior to encapsulation into the IPNs. Cells were then encapsulated into IPNs without doxycycline. Cells were imaged with a Nikon spinning disk confocal microscope under 20x N/A0.75 objective with 445nm laser excitation and the emission filters 511/20 nm for donor (cyan) and 534/30 nm for acceptor-FRET, with 2×2 binning of the Hamamatsu Orca Flash4.0 sCMOS camera at 1 second exposure time per image. Z-stacks were taken every 20 min in the donor and acceptor channels of the FRET biosensor, together with Alexafluor-647 conjugated laminin antibody to label the basement membrane.

Post-processing of biosensor FRET images were performed as previously described (pubmed.ncbi.nlm.nih.gov/23931524/). Briefly, ratiometric biosensor analysis consisted of flatfield correction, background subtraction, image registration, and ratio calculations. Flatfield correction was performed based on the acquisition of a set of Z-stacks of shading images that consisted of cell-free fields of views with identical exposure and field illumination as the biosensor image sets. The donor and acceptor-FRET images were divided by the shading images to achieve the flatfield-correction. A small region of interest in the cell-free background area was selected in the flat-field corrected image sets, and the mean gray value from such a region of interest was subtracted from the whole field of view at each time point at each Z-position to obtain the background-subtracted image sets. The resulting background subtracted Z-stacks of images were segmented by producing and applying binary masks using manual histogram thresholding. The segmented acceptor-FRET channel image stacks were then divided by the segmented donor image stacks to produce the ratio data stacks, followed by an application of a linear pseudocolor lookup table to depict the relative differences in FRET / donor ratio and the corresponding RhoA activity in cells.

### Western Blotting

Cells were lysed on ice for 30 min in a buffer containing 1% NP-40, 50 mM Tris pH 7.4, 150 mM NaCl, 10 mM ethylenediaminetetraacetic acid (EDTA), 1 mM Pefabloc SC (Sigma), and protease inhibitor cocktail (Sigma). The lysate was clarified by centrifugation at 22,000 rcf for 15 min at 4°C. Lysates were resolved by 10-12 % sodium dodecyl sulfate-polyacrylamide gel electrophoresis (SDS-PAGE). Proteins were transferred to polyvinylidene fluoride membranes. After blocking for at least 1 hour in 5% BSA in Tris-buffered saline containing 1% Tween-20 (TBS-T), membranes were incubated with primary antibodies at 1:500 dilution overnight at 4°C. Membranes were incubated with secondary fluorescently labeled antibodies (LI-COR Biosciences, Lincoln, NE, USA) at 1:20,000 dilution for 1 h at RT. Immunoblots were visualized using the Odyssey Imager (LI-COR Biosciences).

### GST-RBD pull-down experiments

HEK293T cells were plated overnight at a density of 0.9 × 10^6^ cells on poly-L-lysine-coated six-well plates. The mVenus-RhoA Q63L and T19N mutants and the RhoA biosensor containing Q63L mutation, and that which contained in addition a p21-binding domain from the p21-activated kinase 1 (PBD), were transfected using the polyethyleneimine (PEI) reagent at 2 µg DNA to 8 µL PEI ratio for each well, according to a published protocol (DOI 10.1002/sita.200500073). After 24 h, cells were lysed in a buffer containing 1% NP-40, 50 mM Tris HCl, pH 7.4, 500 mM NaCl, 10 mM MgCl_2_, 1 mM PMSF, and a protease inhibitor cocktail (Sigma-Aldrich). Lysates were clarified by centrifugation at 22,000 rcf for 15 min at 4°C. Biosensor pull-downs were performed using purified Rhotekin-RBD-agarose beads, as previously described (pubmed.ncbi.nlm.nih.gov/24504947/). To prepare the glutathione (GSH)-agarose beads, 72 mg of GSH-agarose (Sigma-Aldrich) was suspended in 10 ml sterile water and incubated at 4°C for 1 h. The suspension was centrifuged at 1200 rcf for 30 sec at 4°C, and the pellet was washed three times with sterile water, followed by washing two times in a resuspension buffer containing 50 mM Tris, pH 8.0, 40 mM EDTA, and 25% sucrose. To generate GST-Rhotekin-RBD, pGEX-RBD (a gift from G. Bokoch) was transformed into BL21(DE3)-competent strain of bacteria (Agilent Technologies) and propagated in a shaker incubator at 225 rpm and 37°C until an optical density of 0.9 at 600 nm was reached. 0.2 mM of Isopropyl β-d-1-thiogalactopyranoside (IPTG) was added to induce the protein synthesis, at 225 rpm and 24°C overnight. Following an overnight incubation, bacteria were pelleted and resuspended in 20 ml of the resuspension buffer containing 1 mM PMSF, protease inhibitor cocktail (Sigma-Aldrich), and 2 mM β–mercaptoethanol, and rotated for 20 min at 4°C. Following the incubation, 8 ml of the detergent buffer (50 mM Tris, pH 8.0, 100 mM MgCl_2_, and 0.2% [wt/vol] Triton X-100) was added, and the mixture was incubated at 4°C for 10 min on a rotator. Ultrasonication was used to disrupt the cell membrane (4× cycles of 30-s ultrasonication followed by 1 min rest on ice) and centrifuged at 22,000 rcf for 45 min at 4°C. 1 ml of the hydrated and washed GSH-agarose beads were added to the clarified supernatant and incubated at 4°C for 1 h on a rotator. The beads were pelleted by a centrifugation at 1200 rcf for 30 sec and washed four times with the wash buffer containing 50 mM Tris, pH 7.6, 50 mM NaCl, and 5 mM MgCl_2_, followed by resuspension in 500 µl of 50:50 glycerol/wash buffer. Cell lysates were incubated with Rhotekin-RBD–conjugated agarose beads for 1 h at 4°C on a rotator, washed 3x in the lysis buffer, resuspended in the final sample buffer, boiled at 99°C for 5 min, and analyzed by western blotting.

### Antibodies for Western blotting

EGFP (Roche; 11814460001; Clones 7.1 and 13.1; mixture mouse monoclonal), Rac1 (Cytoskeleton.com; ARC03; mouse monoclonal), Cdc42 (Santa Cruz Biotechnology; sc-8401; Clone B-8; mouse monoclonal), RhoA (Santa Cruz Biotechnology; sc-418; Clone 26C4; mouse monoclonal), and β-Actin (Santa Cruz Biotechnology; sc-69879; Clone AC-15; mouse monoclonal). Unless otherwise stated, all primary antibodies were used at 1:500 dilution for Western blots.

### Quantifying deformations of the basement membrane before invasion

To quantify deformations of basement membrane (BM) before invasion, we performed fast-iterative digital image correlation (FIDIC)^61^ with a subset size of 16×16 and spacing of 4 pixels (∼2 to 2.8 µm) on confocal images of the BM before invasion. For each invasive acini, the time and the confocal image of BM at which the invasion occurred was identified. At this invasive plane, confocal images of the BM corresponding to at least 30 minutes and at most 4 hours before the invasion were considered, and FIDIC was performed between two consecutive time-lapse images. To avoid deformations with noisy correlations from FIDIC, we only considered deformations with correlation coefficients above 0.2 (which is at least an order of magnitude greater than the correlation coefficient of noise). To measure radial and tangential deformations in the BM, we transformed into polar coordinates using the center of the BM as origin.

### Modeling and Finite Element simulation

For tumor spheroid, we adopted the multiplicative decomposition of the deformation gradient ***F*** into an elastic tensor ***F_c_*** and a growth tensor ***F_g_***, such that ***F*** = ***F_e_F_g_***. The growth tensor can be expressed by ***F_g_*** = *λ*_g_***I***, where *λ_g_* is the growth stretch and ***I*** is the second order identity tensor. With this definition of the growth tensor, we can then determine the elastic component of the deformation gradient via 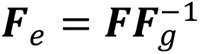. The mechanical behavior of the tumor spheroid may then be described by a Neo-Hookean hyperelastic formulation, with a Cauchy stress given by:

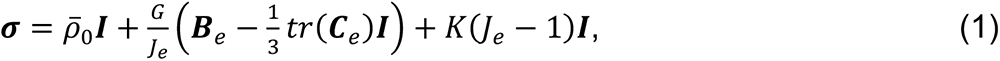

where 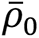 is the contractility of actomyosin, *J_e_* is the determinant of the elastic component of the deformation gradient, 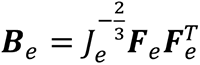 and 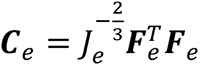 are the left and right Cauchy-Green tensors, respectively, *G* is the material shear modulus, and *K* is the material bulk modulus. In our simulations, BM and hydrogel matrix are linear elastic. While the BM is known to be nonlinear elastic, and the IPNs are viscoelastic and viscoplastic, this simplification should not impact the fundamental physics of this problem. We solve our system using multi-physics software COMSOL to simulate the stress field in BM with the results shown in Fig. 7c.

### Statistical analysis

Most measurements were performed on 2−3 biological replicates from separate experiments. Sample size and statistic test performed for each experiment are indicated in the respective figure legends. Statistical analyses were performed using GraphPad Prism (GraphPad Prism version 9.1.0 for Mac laptop, GraphPad Software, http://www.graphpad.com)

## Acknowledgements

We gratefully acknowledge an Advancing Science in America (ARCS) fellowship for J.C., a National Defense Science and Engineering Graduate fellowship for J.C., a National Science Foundation Graduate Research fellowship for J.C., and a National Institutes of Health National Cancer Institute Grant (R37 CA214136) for O.C. and National Institute of General Medicine Grant (R35 GM136226) for L.H. L.H. is an Irma T. Hirschl Career Scientist. The funding from the Office of Research and Development in the Palo Alto VA Medical Center pays the salary for M. P. M. We thank Professor Jacob Notbohm (University of Wisconin-Madison) for providing the Matlab program to compute curvature of BM.

## Contributions

Conceived and designed the experiments: J.C. and O.C. Performed the experiments: J.C., A.S., S.V., K.L., C.S., and L.H. Contributed analytical guidance: A.S., S.S., L.H., and M.P.M. Z.S. and V.S. designed and implemented the computational model.

Supervised and administered the project: M.C.B., R.B.W. and O.C. Acquired funding: O.C. Wrote the manuscript: J.C., A.S., Z.S., S.V., V.S., and O.C.

## Supplemental Information

**Figure S1.**
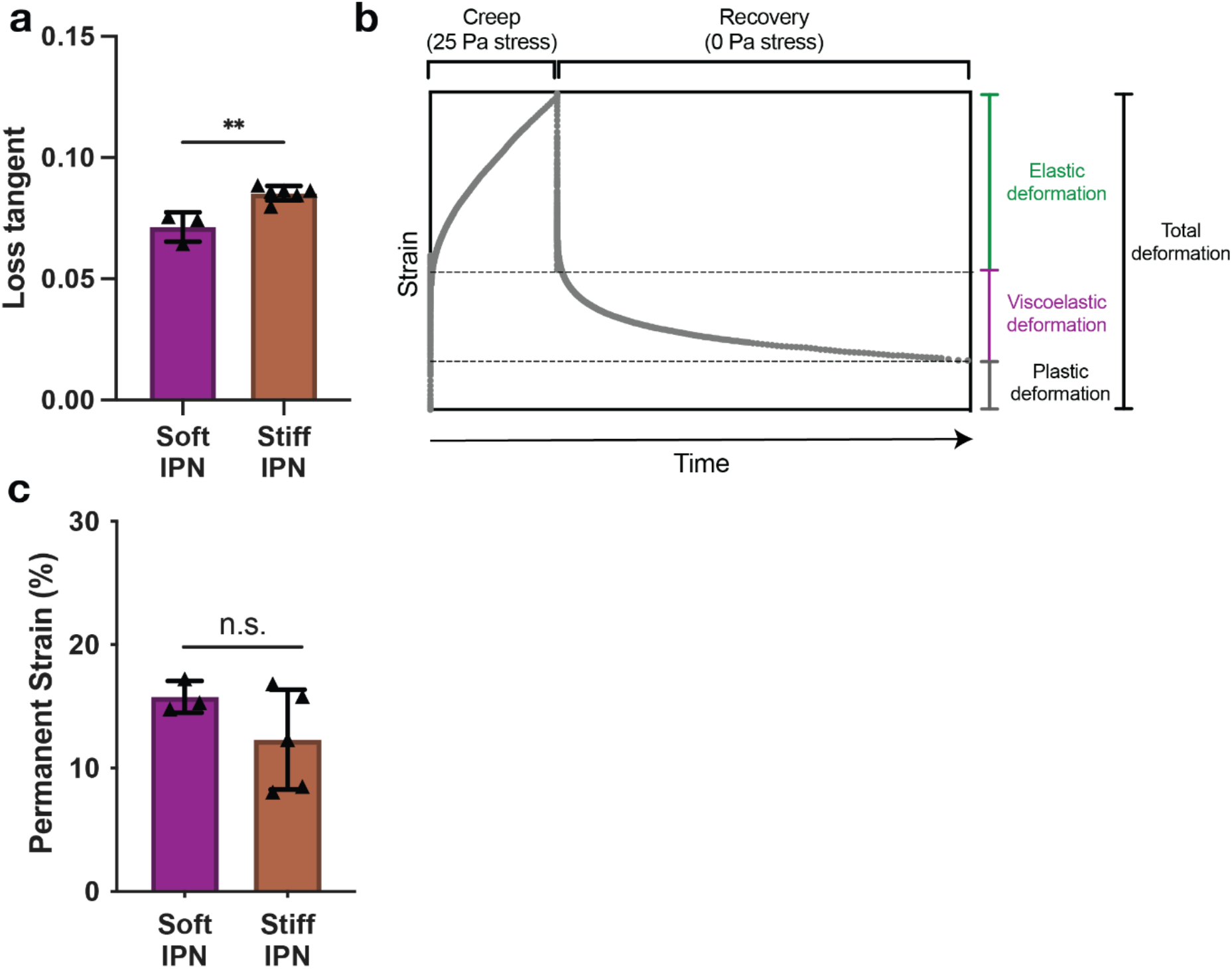
Loss tangent and permanent strain of soft and stiff IPNs. **a,** Loss tangent of alginate-rBM IPNs (soft and stiff). \*\**p* < 0.01, unpaired t-test, *n =* 3 – 6 hydrogels. **b**, Representative creep and recovery test. Example creep and recovery test used to measure plasticity, in which plasticity is quantified as the remaining strain after 10,000 seconds divided by the total deformation during the creep test. **c,** Permanent strain of alginate-rBM IPNs (soft and stiff). ns not significant, unpaired t-test, *n =* 3 – 6 hydrogels.

**Figure S2.**
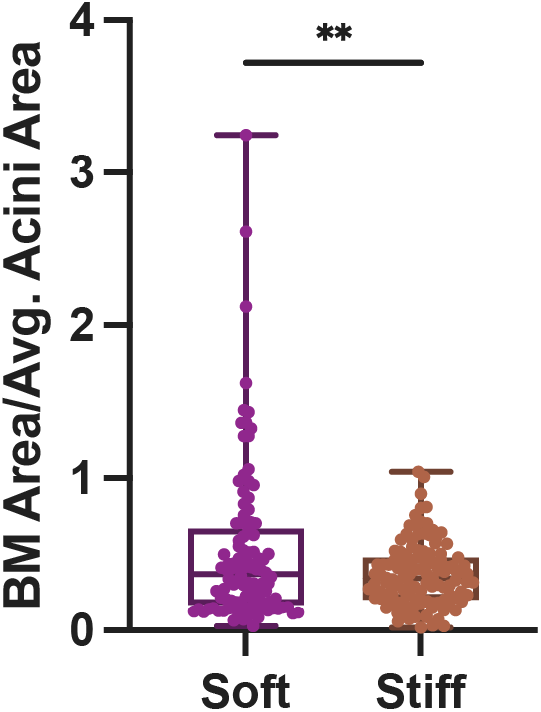
BM metric in soft and stiff IPNs. Quantification of BM area/average acini area in soft and stiff IPNs. \*\**p <* 0.01, Welch’s t-test.

**Figure S3.**
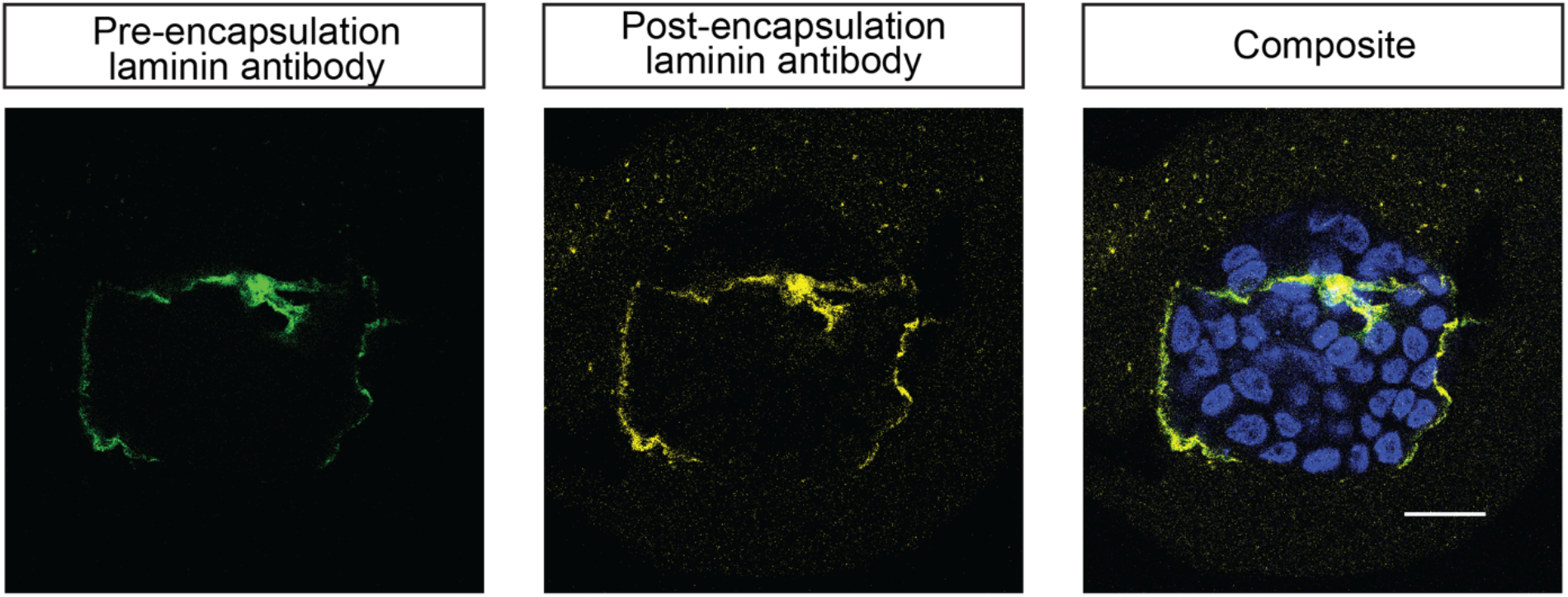
No additional laminin secretion observed in BM after encapsulation of acini into IPNs. Laminin-antibody added to MCF10A acini before IPN encapsulation (green) and laminin-antibody added after IPN encapsulation (yellow) show no new secretion of laminin within 24 hours. Scale bar = 20 μm.

**Figure S4.**
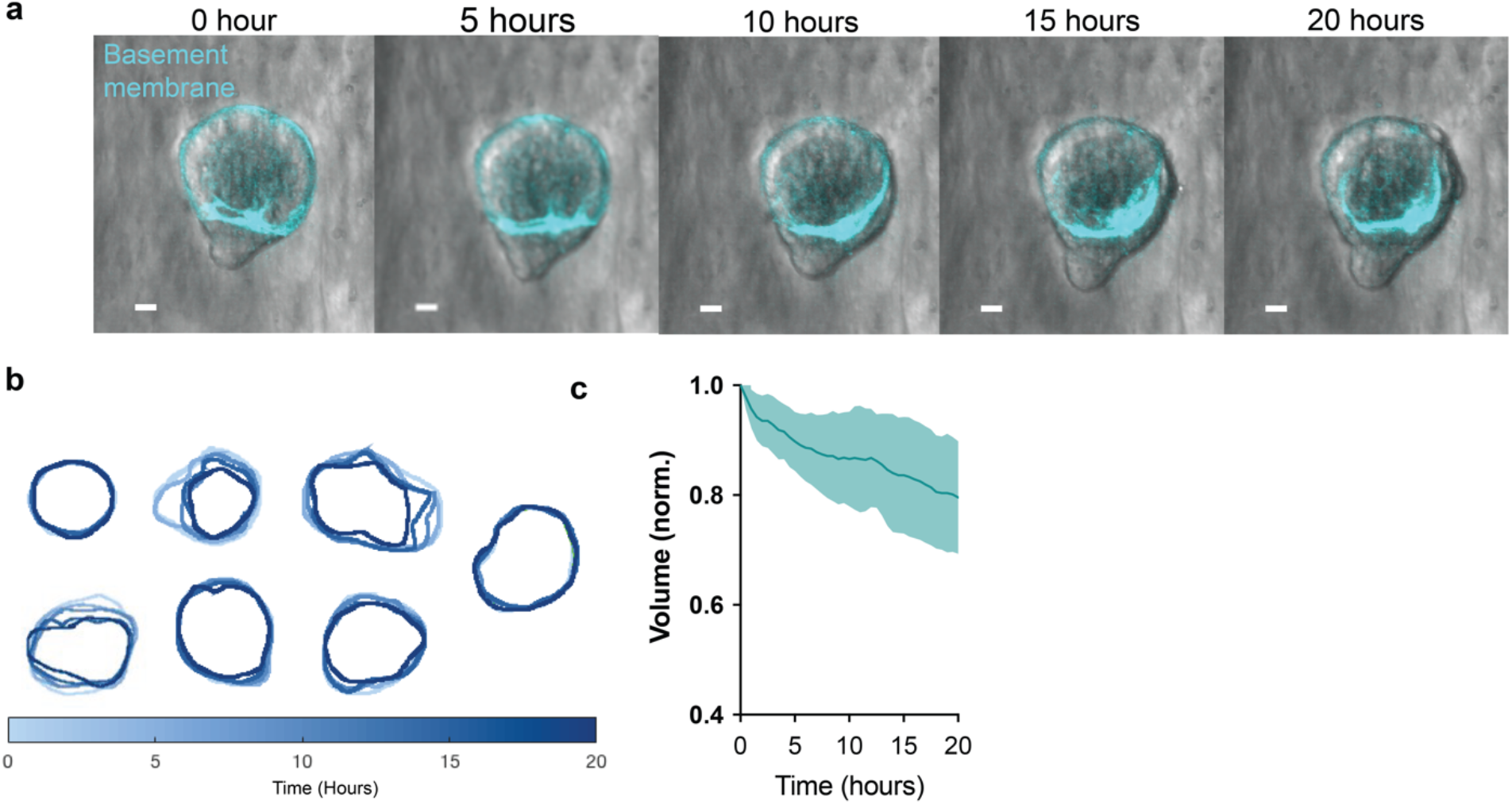
Analysis of collective cell movement in relation to site of BM breaching. **a,** Time-lapse of cell invasion (brightfield) and BM (cyan) movement. Scale bars, 20 μm. **b**, Time-lapse outlines of BM outlines over twenty hours. **c,** Normalized volumes of BM shells over twenty hours.

**Figure S5.**
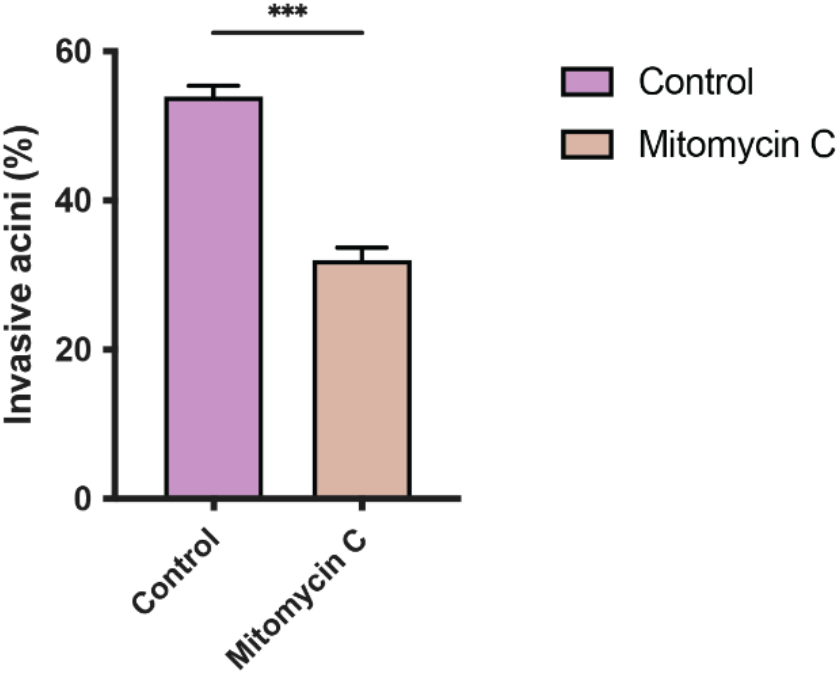
Proliferation contributes to collective cell invasion in stiff IPNs. Quantification invasive acini in stiff IPNs with and without inhibition of proliferation by mitomycin C. \*\*\**p <* 0.001, Unpaired t-test.

**Figure S6.**
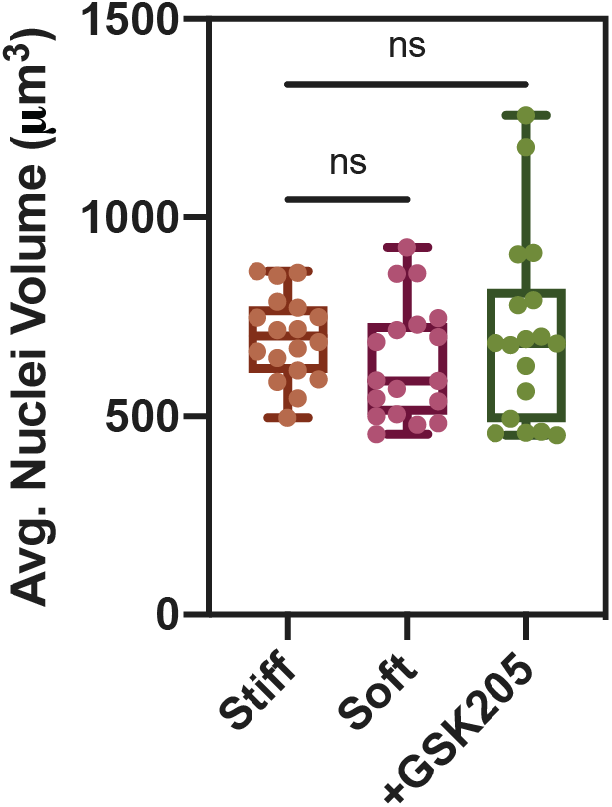
No differences in nuclei volume between soft, stiff, and GSK205 treated stiff IPNs. No differences observed in nuclei volume between soft, stiff, and GSK205 treated stiff IPNs. ns = not significant, ANOVA.

**Figure S7.**
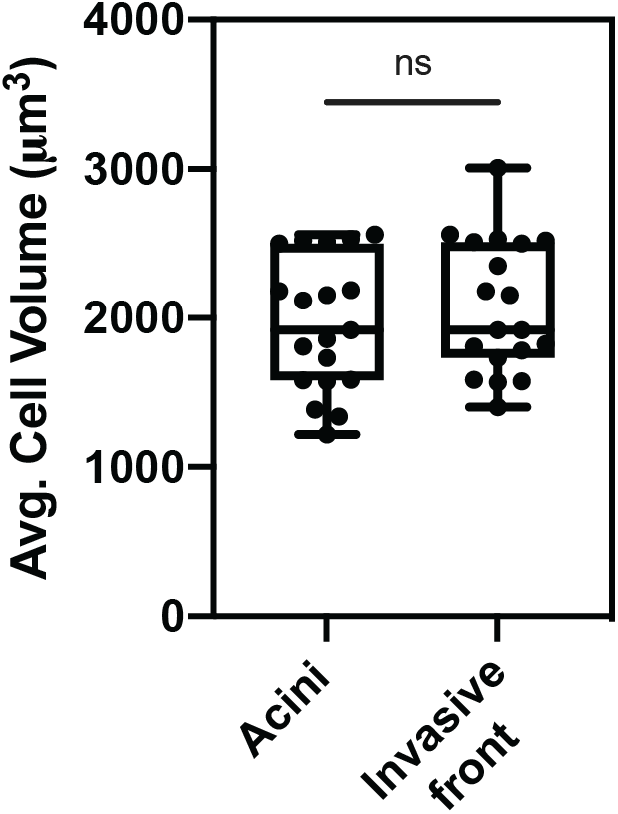
Cell volume in invasive front and acini of invasive acini. No differences in average cell volume between cells in invasive front and cells within the acini of invasive acini in stiff IPNs. ns = not significant, ANOVA.

**Figure S8.**
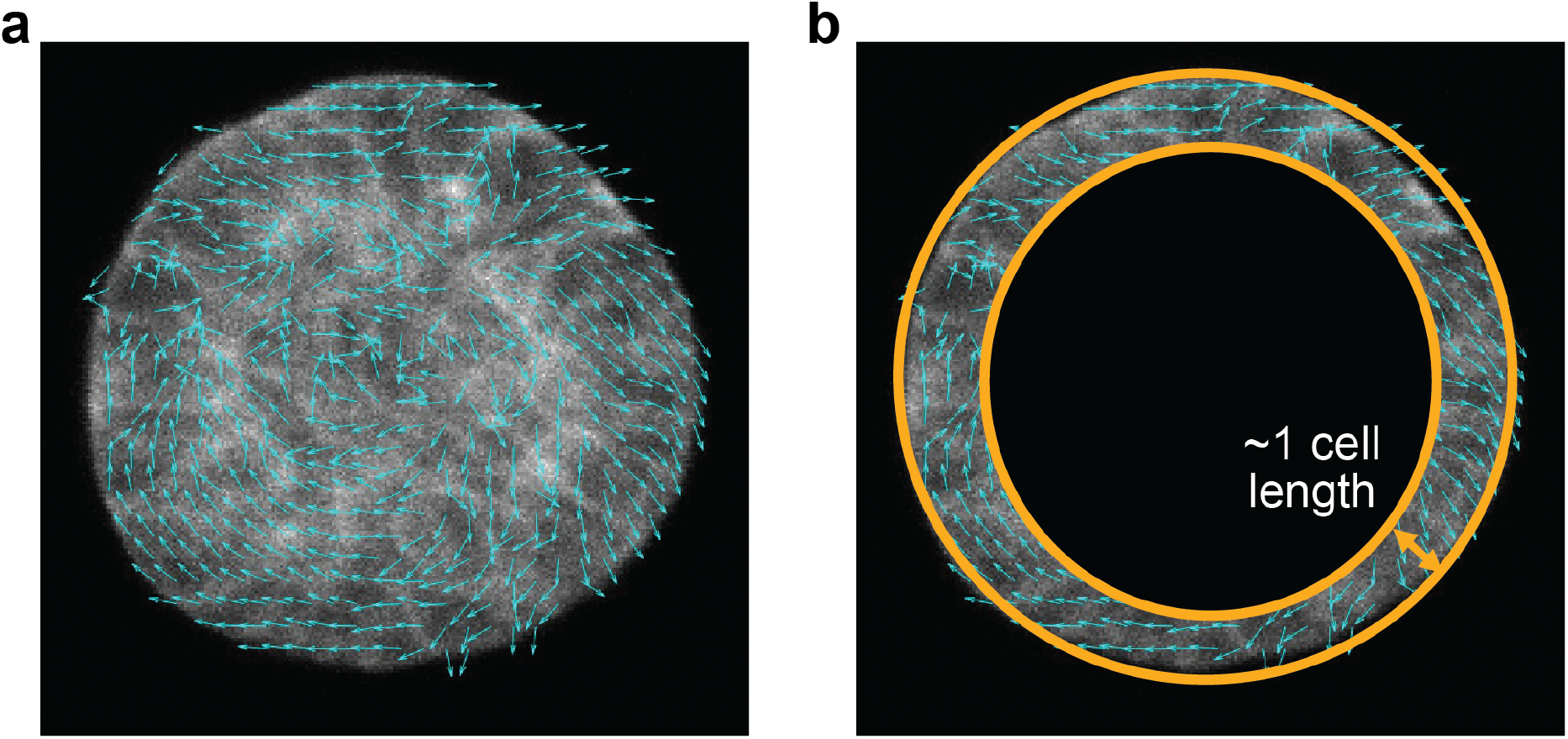
Cells in acini with breached BM collectively exhibit greater radial velocities and decreased tangential velocities compared to cells in acini with intact BM. **a,** Acini actin with cell velocity vectors shown in cyan. **b,** Schematic of region analyzed for cell velocity measurements.

**Figure S9.**
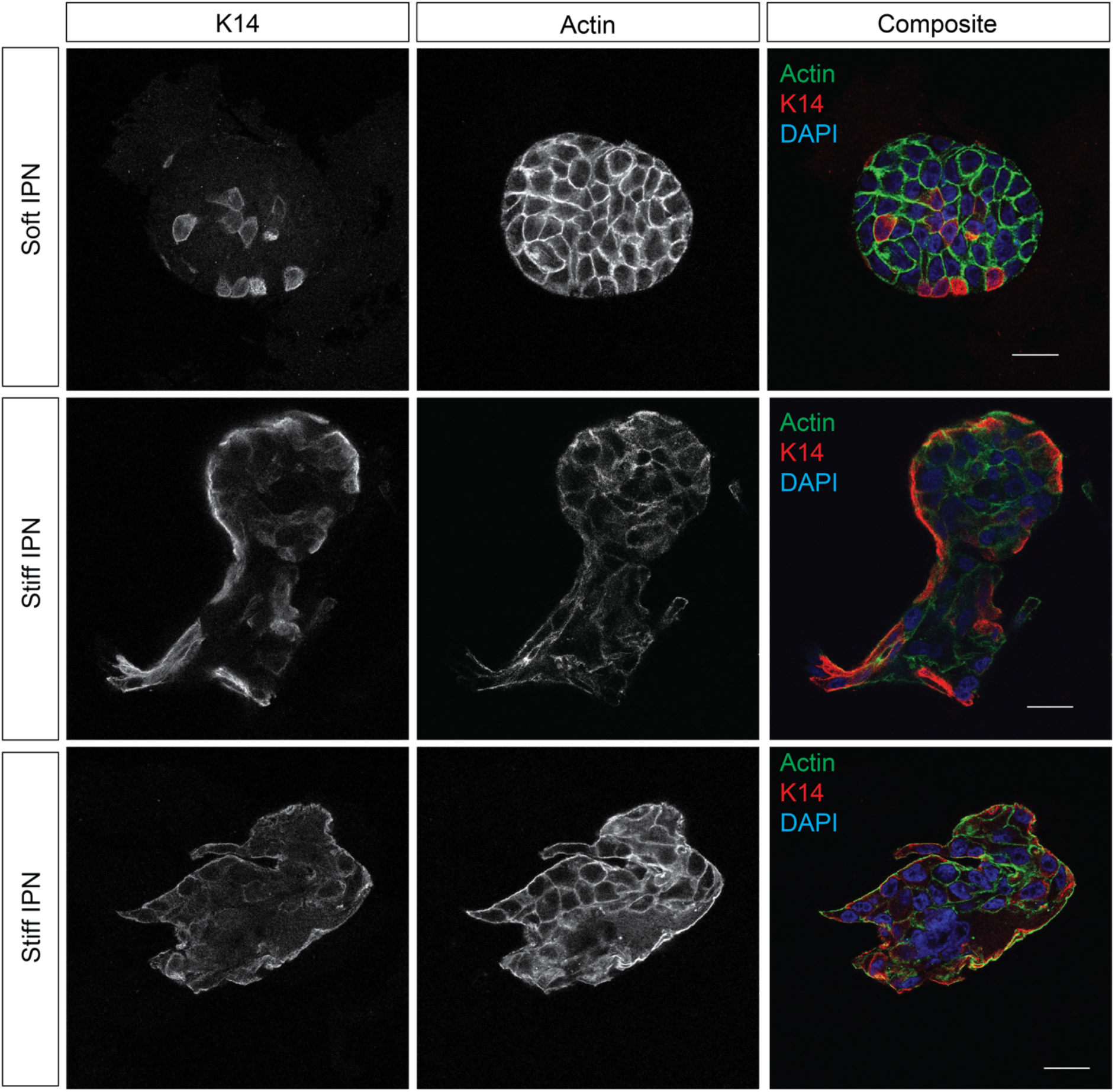
Heterogeneous K14 staining for cells in invasive clusters. Cytokeratin-14 (red), actin (green) and DAPI (blue) staining in acini of soft and stiff IPNs. Scale bar = 25 μm.

**Figure S10.**
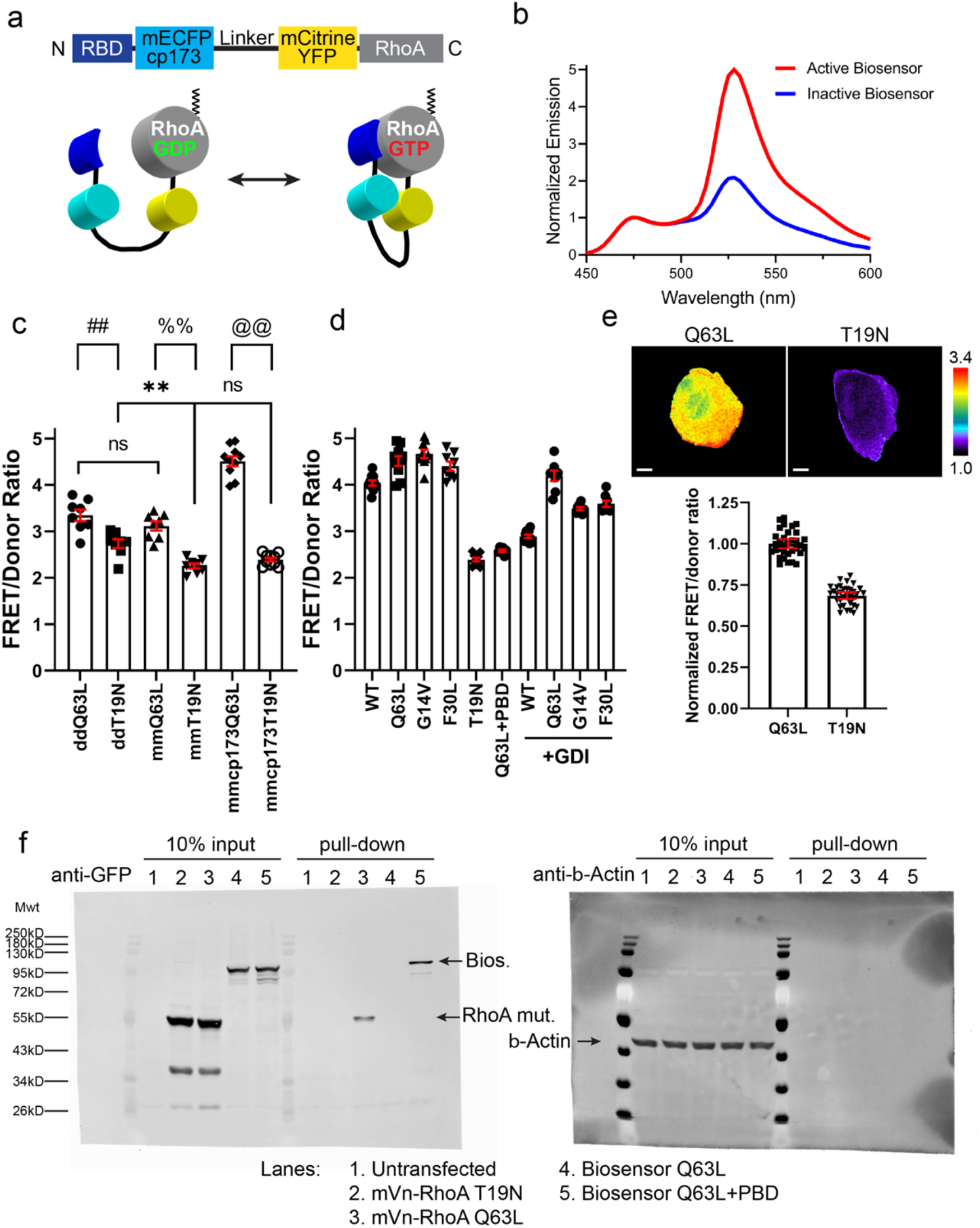
Optimized FRET biosensor for RhoA GTPase. **a**, Schematic representation of the optimized, single-chain, genetically-encoded FRET biosensor for RhoA GTPase, based on the previously published design (pubmed.ncbi.nlm.nih.gov/16547516/). Fluorescent proteins are replaced with monomeric versions containing the A206K mutations, and the dipole coupling angle is optimized by using a circular permutation of the donor at position 173. **b**, Representative, normalized fluorescence emission spectra are shown, from the constitutively activated Q63L (“Active Biosensor”) and the dominant negative T19N (“Inactive Biosensor”) mutant versions of the biosensors overexpressed in live cell suspensions of HEK293T cells, excited at 433 nm and the emission scanned from 450 nm to 600 nm. Spectra are normalized to the emission maxima of the donor fluorophore at 427 nm. **c**, Changes in FRET/donor ratio responses comparing active versus inactive mutant versions of the RhoA biosensor during optimization of the fluorescent protein FRET pair. “dd” indicates original RhoA biosensor (pubmed.ncbi.nlm.nih.gov/16547516/) containing the dimerizing A206 residue in both the donor and the acceptor fluorescent proteins, “mm” indicates A206K monomeric mutations are introduced into the original ECFP and the Citrine YFP, and “mmcp173” indicates A206K monomeric mutations in both the donor and acceptor fluorescent proteins in addition to circular permutation of the donor at amino acid position 173. Student’s t-test, two-tail analysis: ## p= 0.001339; %% p = 1.8517×10^-6^; @@ p = 2.1825×10^-13^; ** p= 0.0005084; ns (between ddQ63L versus mmQ63L mutants) p=0.159136; ns (between mmT19N versus mmcp173T19N) p = 0.05596; n = 7 experiments for dd and mm, and n = 10 for mmcp173, shown with SEM. **d**, Fluorometric FRET/donor emission ratio of the optimized RhoA biosensor overexpressed in HEK293T cells. WT biosensor expression and the Q63L, G14V, and F30L constitutively activated mutant biosensors showed high emission ratios. The dominant negative T19N mutant biosensor, constitutively activated (Q63L) biosensor with non-specific binding domain (PBD) showed low emission ratios. 2-fold excess expression of the guanine nucleotide dissociation inhibitor-1 (GDI) showed reduced ratio for those that are targeted by GDI, whereas the Q63L mutant that does not bind GDI produced elevated ratio. Student’s t-test, two-tail analysis: WT compared against Q63L, G14V, F30L, and T19N p=0.0008309, 1.4319×10^-5^, 0.004302, and 4.8370×10^-15^, respectively; Q63L compared against Q63L+PBD, p = 1.1849×10^-9^; and +GDI against respective conditions without GDI, WT p = 3.3425×10^-13^, Q63L p = 0.05522, G14V p = 7.0422×10^-7^, and F30L p = 9.4961×10^-6^. N = 10 experiments for WT, Q63L and T19N, N = 9 for G14V, N=8 for F30L, and all +GDI conditions, N = 6 for Q63L+PBD; all shown with SEM. **e**, Ratiometric microscopy measurements of the transiently expressed RhoA biosensor mutants in MCF10A cells under 60x magnification, and the quantification of the whole-cell average ratio values. White bars = 10 µm. Linear pseudocolor corresponds to the indicated scaling limits. Student’s t-test, two-tail analysis, p = 3.6545×10^-5^, N = 3 experiments, shown with 95% confidence intervals of the aggregate data points from the pooled datapoints. **f**, Competitive pull-downs of the optimized RhoA biosensor. Lane designations are also shown. The Q63L constitutively activated RhoA biosensor is pulled down by excess exogenous GST-RBD, only when the built-in RBD domain of the biosensor is exchanged for a non-Rho-specific PBD.

**Figure S11.**
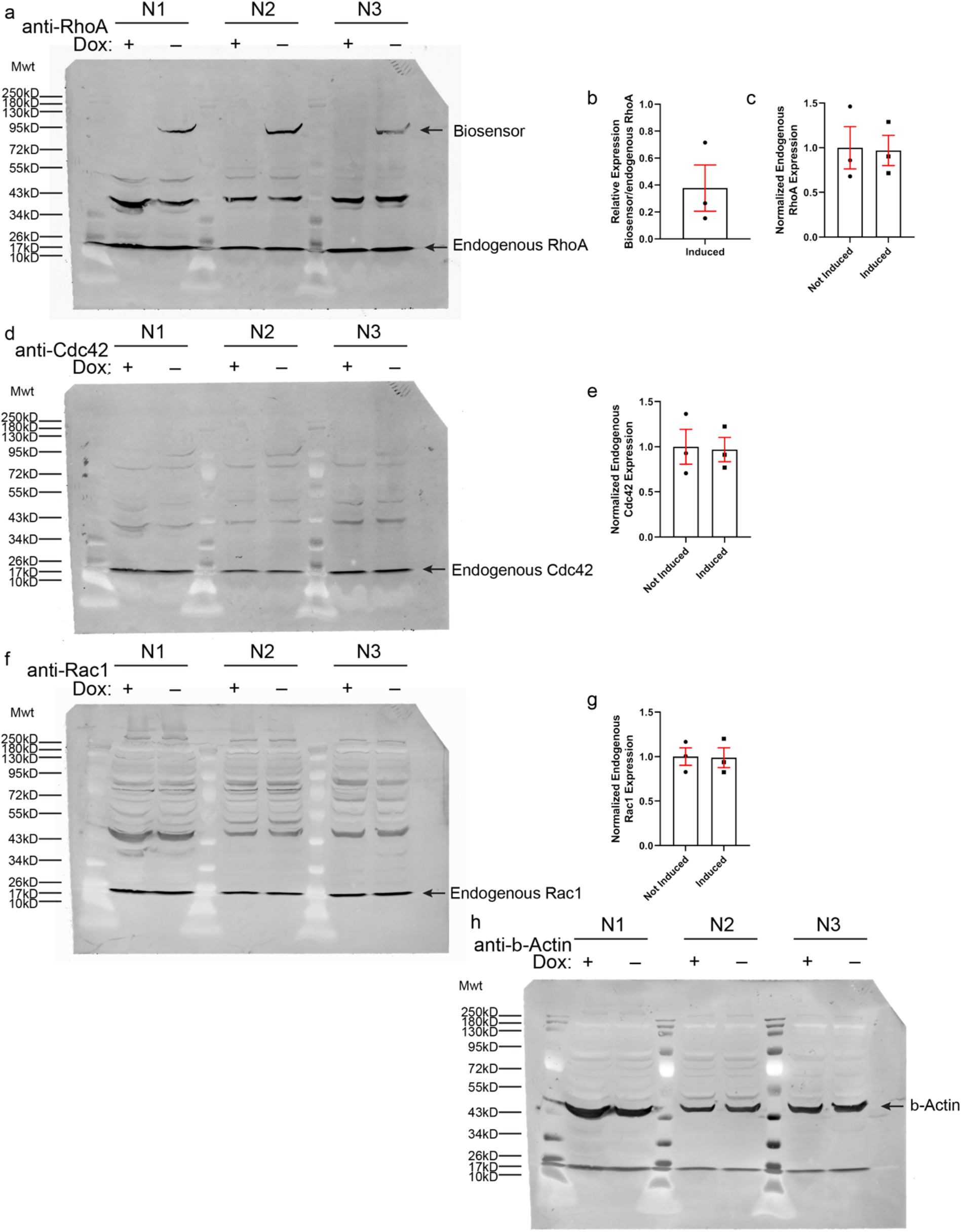
Biosensor expression analysis in MCF10A cells. **a**, Inducible expression of the optimized RhoA biosensor in MCF10A cells, stably incorporating the biosensor under the tetracycline-OFF regulation. MCF10A cells stably transduced with the tet-OFF inducible biosensor expression system were FACS sorted to enrich for the biosensor-positive cell population and analyzed following the same biosensor induction protocol used in the current biological assays. **b**, Quantification of (**a**); induced RhoA biosensor band intensities compared to the endogenous RhoA, indicating 37.8% +/-17.2% of the endogenous RhoA levels, shown with SEM, N = 3 experiments. **c**, Quantification of (**a**); induced expression of the RhoA biosensor does not affect the relative expression levels of endogenous RhoA. Student’s t-test, two-tail, paired-analysis. p = 0.7093, N = 3 experiments. **d**, Western blot detection of endogenous Cdc42, with or without RhoA biosensor induction. **e**, Quantification of (**d**); induced expression of the RhoA biosensor does not affect the relative expression levels of endogenous Cdc42. Student’s t-test, two-tail, paired-analysis. p = 0.650, N = 3 experiments. **f**, Western blot detection of endogenous Rac1, with or without RhoA biosensor induction. **g**, Quantification of (**f**); induced expression of the RhoA biosensor does not affect the relative expression levels of endogenous Rac1. Student’s t-test, two-tail, paired-analysis. p = 0.7294, N = 3 experiments. **h**, β-Actin loading control for the Western blots in (**a**), (**d**), and (**f**).

**Table S1.** Material parameters for modeling.

| Parameters | Meaning | Value | Reference |
| --- | --- | --- | --- |
| $r_t$ | Tumor radius | $50\ \mu m$ | This paper |
| $G_m$ | Shear modulus of gel | 3000 Pa (stiff IPN) | This paper |
| $G_{BM}$ | Shear modulus of BM | 8.2 kPa | Ref. 46 |
| $t_{BM}$ | Thickness of BM | $1\ \mu m$ | This paper |
| $\lambda_g$ | Growth factor | 2.5 | This paper |
| $\rho_0$ | Contractility of actomyosin | 700 Pa (stiff IPN) | Ref. 62 and this paper |
| $\rho$ | Exerted contractility of actomyosin | 1400 Pa (stiff IPN) | This paper |
| $k_{cell}$ | Bulk modulus of tumor | 800 Pa | Ref. 62 |

